# Collinearity of Decomposed Energy Terms in MM-P(G)BSA Binding Free Energy Calculations of Protein-Protein Complexes

**DOI:** 10.64898/2026.06.24.734195

**Authors:** Azize Sevim, Abdulkadir Kocak

## Abstract

The molecular mechanics (MM) Poisson-Boltzmann (PB) or generalized Born (GB) surface area methods (MM-P(G)BSA) are among the most commonly used end state approaches for the calculation of the binding free energies (BFEs) in computational drug design and screening studies. Their thermodynamic cycle is based on the decomposition of the free energy change into several energy terms, including molecular mechanics (MM) electrostatic and van der Waals terms representing the gas phase binding energy change, as well as implicit solvation energies comprising polar solvation, calculated by solving PB equation or its reduced form, GB, and nonpolar solvation surface area (SA) terms. Although these terms are additive, and thus the sum of these components should yield the total free energy change, most of these terms are represented with coefficients that are physically meaningful but empirical. To date, a great deal of effort has been made to improve the accuracy of these methods in the prediction of experimental binding free energies, at least relatively, i.e., relative binding free energy (RBFE), resulting in totally empirical coefficients on energy terms and breaking the underlying physics. Furthermore, although these methods originated from protein-ligand RBFE estimation, there have been examples or even tutorials using these methods for protein-peptide or protein-protein interactions (PPIs), due to the same simple physics of binding at the bound state, without sufficient hesitation regarding their accuracy limitations. Here, we thoroughly evaluate MMP(G)BSA methods in all variants available in the literature for a diverse protein-protein complex set and reveal their true RBFE accuracy by means of Pearson correlation with experimental values, with at most R = 0.42. We also demonstrate that the assumption of independent fitting coefficients for decomposed energy terms could not only break the physics but also statistically represent overfitting. Through analytic derivation and large-scale molecular dynamics simulations, we show that (i) the protein-ligand (PL), protein-peptide or protein-protein (PP) Coulomb interaction energy and the GB solvation correction are almost perfectly collinear (R^2^≥0.99) reflecting their designed role as vacuum electrostatics plus solvent screening, and (ii) the van der Waals interaction and SA term also exhibit strong correlation. Interaction entropy (IE) and C2 entropy corrections, which are also found to be strongly dependent on each other, worsen the overall accuracy of the methods due to the large energetic fluctuations of these extended protein-protein systems. The findings of collinearity hold both at the level of instantaneous trajectory fluctuations and when averaged across a diverse set of 138 PP complexes and persist in both single-trajectory and three-trajectory MM-GBSA protocols. Our results show the limitation of the methods for protein-protein complexes and caution against using decomposed MM-GBSA terms as independent predictors in regression models and suggest instead combining correlated terms into effective polar, nonpolar, and entropic contributions, while noting the adverse impact of these entropy terms.

## 1. Introduction

In computer aided drug design (CADD), binding affinity is a fundamental quantity for understanding thermodynamics and thus drug-like features of ligands. Although there are numerous methods and strategies, end-state approximate methods such as Molecular Mechanics (MM) Poisson-Boltzmann Surface Area (PBSA) and Generalized-Born Surface Area (GBSA) are still widely used due to their simplicity and relatively low computational cost, enabling their utilization for high-throughput virtual screening (HTVS). These methods just rely on bound and unbound states of the ligands to their biological relevant targets. Rather than absolute binding free energies (ABFE), these approximate methods aim to find relative binding free energies (RBFE) among different ligands to a common target or the same ligand to different targets such as variants of a protein. MM-PBSA (or GBSA) are applied as post-molecular dynamics (MD) or also used as refining docking scores [1].

Binding free energy Δ*G*_*bind*_ is a fundamental thermodynamic quantity that quantifies the strength of the association between a protein (receptor) and its partner (ligand, peptide or another protein) [2,3]. It represents the change in Gibbs free energy between the bound complex and unbound states of components (receptor, ligand, peptide, etc.) in solution [4].

Molecular Mechanics Poisson-Boltzmann Surface Area (MM-PBSA) and Molecular Mechanics Generalized-Born Surface Area (MM-GBSA) approaches based on molecular dynamics (MD) simulations are popular end-point methods used to estimate binding free energies with a balance between computational efficiency and accuracy [1,5–11]. Unlike rigorous alchemical methods that require sampling multiple non-physical intermediate states, these methods evaluate only the bound and unbound states of the system [1,2]. The MMP(G)BSA methods are considered more accurate than most docking scoring functions and are widely applied in high throughput virtual screening (HTVS) as well as lead optimization of the drugs [5,12].

Although these methods primarily focused on protein-ligand systems where ligand could be an inhibitor, the governing physics of A- and B-type complexation to form AB also holds for other molecular systems, such as protein-peptide and protein-protein interactions. Thus, program packages like MMPBSA.py from AmberTools or gmx_MMPBSA.py also provide tutorials on how to use the scripts for protein-peptide/protein (PP) complexes, without rigorous validation regarding their limitations for such comparatively larger systems and without clear awareness of accuracy in such extended systems. Indeed, there have been a limited number of studies benchmarking the success of these methods on PP systems.

The general idea in MM-PB(GB)SA is to decompose the total binding free energy (Δ*G*_*bind*_) into several independent physical contributions [1,6,13,14]. The standard equation represents binding as a sum of four major components [15]. The molecular mechanics (MM) term accounts for non-covalent interactions including both van der Waals and electrostatic contributions in so-called “gas-phase”, in which the solvent is removed although the conformational space is still sampled in the solvent environment. [1,2,16]. The solvation free energy consists of a polar term (obtained by solving the Poisson-Boltzmann (PB) or Generalized-Born (GB) equations) and a nonpolar term (typically estimated as proportional to the solvent-accessible surface area, SASA or solvent-accessible volume, SAV) [2,12,16]. The entropy term is usually ignored or calculated using harmonic approximation via normal-mode analysis (NMA), quasi-harmonic approximation via principle component analysis (PCA) or via energy fluctuation as interaction entropy (IE) and C2 entropy. [1,5,7,15–17].

One of the main limitations of MM-PB(GB)SA methodology lies in the assumption of decomposed energetic contributions as independent and additive terms, which is not supported by a rigorous assessment of their interdependence. In standard protocols, the binding free energy is typically obtained by directly summing electrostatic, van der Waals, polar solvation, and nonpolar solvation components, supposing that each term represents a separable physical contribution. However, most of these energy terms depend on empirical parameters whose values may vary depending on the selected model and parametrization scheme. For example, the polar solvation contribution in PB(GB) models depends strongly on parameters such as the solute dielectric constant, while the nonpolar solvation term is commonly estimated from the solvent-accessible surface area (SASA) scaled by an empirical surface tension coefficient (γ). Although some of these parameters are based on physical basis, literature frequently reports fitted parameter sets chosen to maximize agreement with experimental binding affinities rather than to preserve strict physical interpretability. For instance, surface tension or pressure coefficients are chosen as 0.005 or 0.0075. Similarly, for protein–ligand electrostatic interactions and polar solvation terms, the internal dielectric constant (ε_in_) is often selected within a range from the vacuum value of 1 to the solvent value of 80, depending on which value provides the best agreement with experimental data. The physical basis for this last example is as follows: Since the number and distribution of acidic/basic amino acids in the protein structure are not equal and homogeneous, the internal dielectric constant will also vary. However, selecting a single dielectric constant that provides the best fit across an entire dataset raises questions regarding the true physical meaning of this parameterization. In addition to the original MM-PB(GB)SA equation, recent studies have shown that other energy terms included in these formulations are also represented using empirical parameters, regardless of whether they have a physical basis. Consequently, the direct summation of these parametrized energy terms may obscure intrinsic coupling and statistical dependencies between energetic contributions. In addition, to improve quantitative agreement with experimental data, researchers often use reparametrized models where regression coefficients are fitted against experimental measurements [2,18–23]. This is conceptually similar to the Linear Interaction Energy (LIE) method, where the binding energy is assumed to be linearly proportional to the differences in van der Waals and electrostatic interactions, scaled by empirical parameters α and β [24]. However, relying on multiple adjustable terms may lead to hidden collinearity, where different energy components are statistically redundant or compensate for each other. For example, ligand interactions with “hot-spot” residues often compensate for the loss in polar interactions with the solvent upon binding, making it difficult to distinguish their individual impacts [25]. Inclusion of extra, suspicious terms in a regression model may also result in overfitting, providing a high correlation on training data that fails during blind prospective testing [25,26]. Overall, although the independence assumption significantly simplifies computational treatment and enhances efficiency, it limits the physical rigor of the model by neglecting the intrinsic coupling between energetic terms. As a result, both the accuracy and transferability of MM-PBSA predictions may be constrained in complex biomolecular systems.

From statistical perspective, the emergence of all these variants of MMP(G)BSA approaches and thus the use of empirical parameters have raised the question of whether these energy components are independent variables and, consequently, whether they can be included in the energy equation with separate empirical parameters. Therefore, collinearity analyses among the energy terms are necessary to determine how many independent variables exist in binding free energy equation.

In this study, we systematically investigate the statistical dependencies underlying MM-PB(GB)SA energy decomposition and their implications for binding free energy (BFE) prediction in PP systems. To this end, MM-PB(GB)SA calculations were performed on a curated dataset of 138 protein–peptide complexes, enabling both trajectory-level and ensemble-level analyses.

## 2. Theory and Background

The Molecular Mechanics (MM) Poisson-Boltzmann Surface Area (PBSA) [1,14,27–31] and Generalized-Born Surface Area (GBSA) [28,32–34] methods are widely used end-state approach for estimating the binding free energies of biomolecular complexes. MM-PB(GB)SA method assumes that the binding free energy can be estimated using only the final states of the system and considers only the bound and unbound states of the system for evaluation.

The general idea in these calculations is to decompose the total binding free energy change (Δ*G*_*bind*_) into several physical contributions [1,6,16]. The standard equation represents binding as a sum of classical molecular mechanics (MM) energy terms, the polar solvation energy calculated using either the Poisson-Boltzmann (PB) or generalized Born (GB) approaches, the nonpolar solvation energy estimated via continuum surface area (SA) models, and the entropy change upon binding (−*T*Δ*S*).

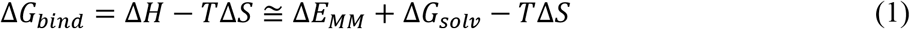

assuming pΔV ≈ 0 for condensed phases and thus Δ*H* = Δ*E*_*MM*_ + pΔV ≈ Δ*E*_*MM*_

where

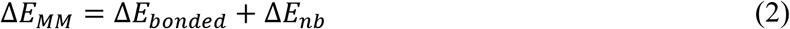

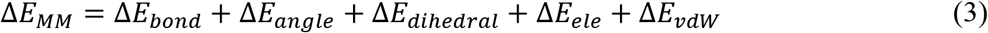

where each term is calculated by means of difference between the complex (AB) and individual (A and B) molecules (i.e., AB-A-B). For non-covalently interacting molecules, bonded terms cancel out and the equation reduces to:

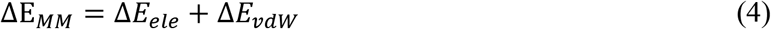

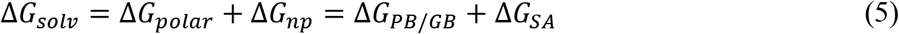

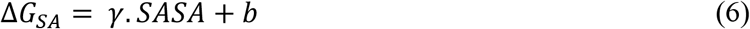

where, Δ*E*_*MM*_, Δ*G*_*solv*_ and −*T*Δ*S* represent the changes in gas-phase molecular mechanics energy, solvation free energy, and entropy changes, respectively, upon binding. These individual energy components are evaluated using an ensemble of conformational snapshots extracted from MD simulations. In principle, the calculation of binding free energy using this approach requires that the above equations be formulated based on three independent simulations: (i) the protein– ligand complex in solvent, (ii) the protein alone in solvent, and (iii) the ligand alone in solvent.

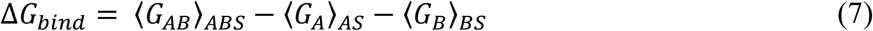

Here, the subscript outside the angle brackets denotes the simulation environment (e.g, ABS=AB complex+solvent system) over which the average is taken, while the subscript inside the brackets indicates the specific energy term being evaluated (e.g., Δ*G*_*AB*_ is the free energy of AB complex). This approach requires three separate (A in solvent, B in solvent, AB in solvent) simulations.

Although the formulation known as 3A-MM-PBSA (three-average), which represents the original implementation of MM-PBSA, fully satisfies the thermodynamic cycle described in **Figure 1**, this method has been shown to give low precision in reproducing experimental values due to phase space overlap problems occurred by three separate simulations. The MM-PBSA thermodynamic cycle estimates the binding free energy in aqueous solution by combining the gas-phase binding energy with solvation effects. Specifically, the total free energy change is obtained by adding the binding energy in vacuum to the difference between the solvation free energy of the complex and the sum of the solvation free energies of the receptor and ligand separately. In practice, all required terms can be derived from a single molecular dynamics trajectory by extracting configurations of the complex, receptor, and ligand, and evaluating their solvation energies using the Poisson–Boltzmann equation. The entropic contribution to binding can then be approximated through normal mode analysis [35].

**Figure 1.**
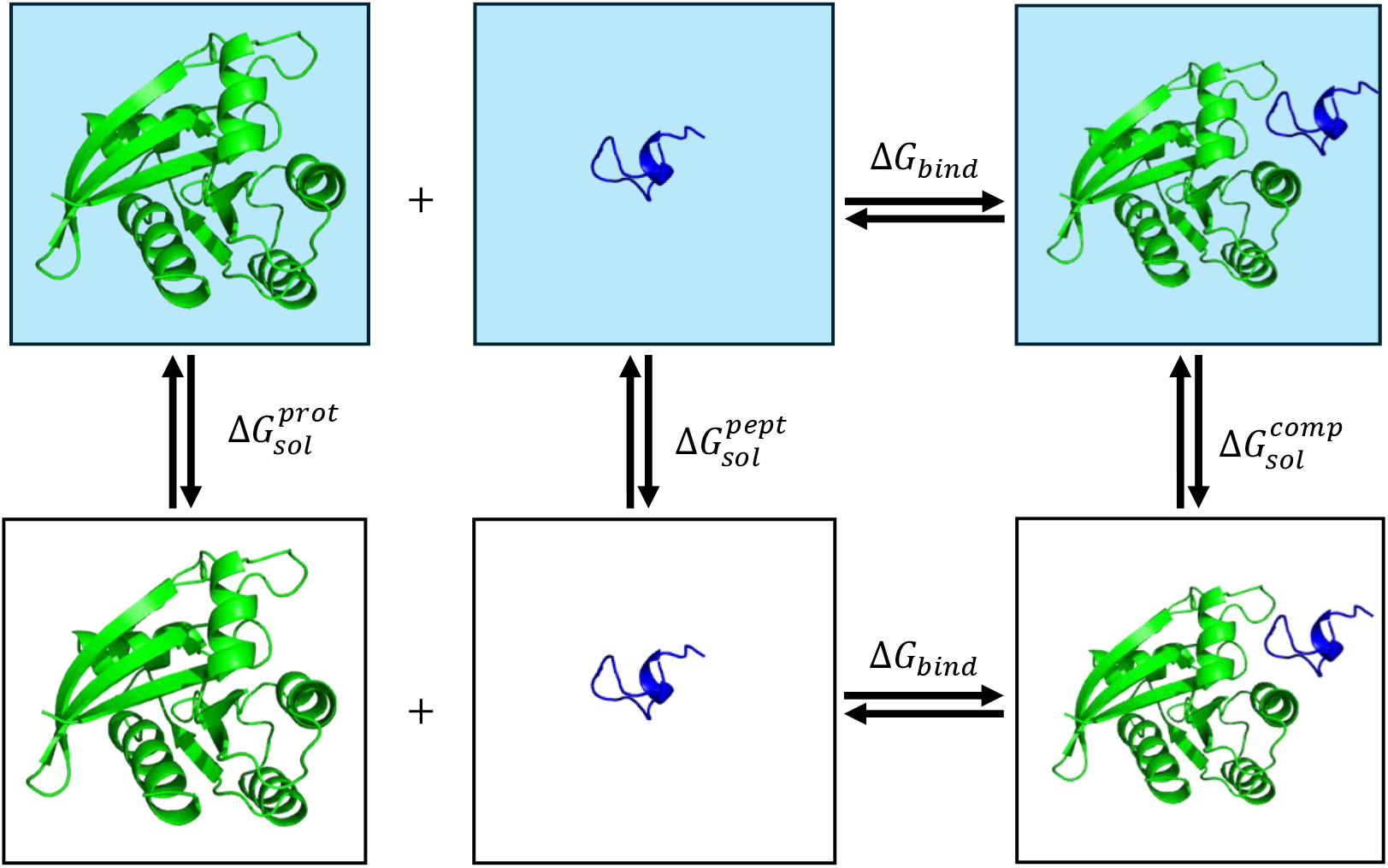
Thermodynamic cycle underlying MM-PB(GB)SA binding free energy calculations. Thermodynamic cycle illustrating the decomposition of binding free energy into molecular mechanics and solvation contributions in MM-PB(GB)SA calculations. The binding free energy is estimated from the free energy difference between the bound complex and the separated receptor and ligand states.

An alternative approach can be employed in which protein and ligand coordinates are directly extracted from a single trajectory generated for the complex. This approach is known as 1A-MM-PBSA (single-trajectory, ST) and incomplete in terms of thermodynamic cycle. Here, the binding free energy in solution is determined as the sum of so-called gas-phase binding energy, corresponding to the A-B interaction energy in the absence of solvent molecules, and the change in solvation free energy upon binding. The gas-phase binding energy is further decomposed into its electrostatic and van der Waals components. The change in solvation free energy (SFE) upon binding is calculated using implicit solvation models. Moreover, SFE is also decomposed into polar and nonpolar contributions. The polar solvation energy is evaluated using either the Poisson-Boltzmann (PB) or Generalized-Born (GB) approaches, whereas the nonpolar solvation energy is estimated as proportional to the solvent-accessible surface area (or volume) of the interacting region with a proportionality constant, surface tension (or pressure) coefficient. The total binding free energy change is calculated according to the following expression:

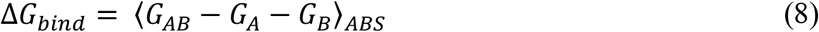

Previous studies have demonstrated that binding energies obtained using this approach tend to yield more precise results [27] since all the components use the same exact trajectory/phase space and the errors cancel out.

However, this approximation can introduce significant errors in free energy calculations because of several factors. For instance, the energy of the system is described at the molecular mechanics (MM) level, and the nature of these terms is rather coarse. In addition, the continuum solvent is only implicitly defined. Furthermore, the variability in the total charge of proteins further limits its applicability. For these reasons and others, this method is not suitable for predicting absolute-binding free energy (ABFE). Owing to its computational efficiency, it is still widely employed, in estimation of relative-binding free energy (RBFE) in computer-aided drug design (CADD) studies.

In Equation 4, the electrostatic energy term (*E*_*ele*_) is typically computed using Coulomb’s law, with atomic partial charges obtained from the molecular mechanics (MM) force field. Vacuum Coulomb between A and B:

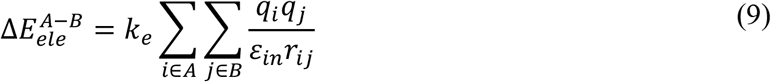

The van der Waals energy term (*E*_*vdW*_) is modeled by Lennard-Jones (LJ) potential function:

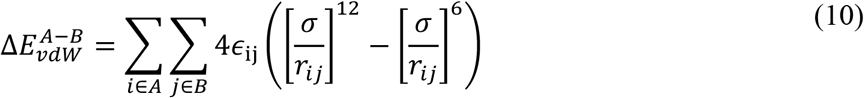

The polar solvation energy (Δ*G*_*polar*_) is calculated by solving the Poisson-Boltzmann equation (PBE):

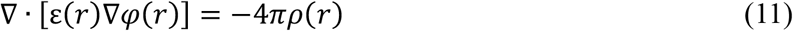

where *φ*(*r*) represents the electrostatic potential of the solute, *ε*(*r*) denotes the spatially dependent dielectric constant, and *ρ*(*r*) corresponds to the fixed charge density. Because the computed polar contribution is sensitive to the selected solute dielectric constant (*ε*_*solute*_), this parameter was systematically varied to evaluate its impact on predicted binding free energies.

MM-PBSA is frequently applied alongside the closely related MM-GBSA method, as both approaches rely on the same input data to estimate binding free energies within a continuum solvent framework. The primary distinction between them lies in how the polar solvation energy is computed. In MM-GBSA, this term is evaluated using the Generalized-Born (GB) model [16], which provides an analytical approximation to the Poisson-Boltzmann equation.

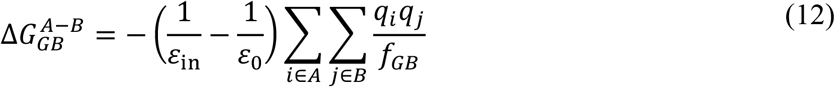

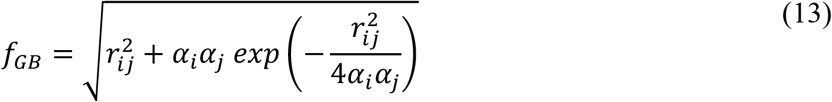

This approximation significantly reduces computational cost, although it may come at the expense of accuracy, depending on the system under study. The GB formulation represents solute atoms as charged spheres characterized by their partial charges (q), internal dielectric constant (*ε*_in_), solvent dielectric constant (*ε*_0_), interatomic distances (*r*_*ij*_), and effective Born radii (α).

The nonpolar contribution to the binding free energy is traditionally approximated as being proportional to the solvent-accessible surface area (SASA) of the solute, expressed in equation 6. In this formulation, γ represents the surface tension coefficient associated with the energetic cost of cavity formation in water, while b is an empirical offset term. The surface tension coefficient and the internal dielectric constant used for the electrostatic terms are typically treated as empirical parameters, chosen to provide the best agreement with experimental data. The internal dielectric constant (*ε*) is most set to 1, corresponding to the vacuum dielectric constant and solvent dielectric constant *ε*_0_ is set to 80.

The entropic contribution can be efficiently estimated using the interaction entropy (IE) approach [36–38]. In this method, the entropy term is expressed as

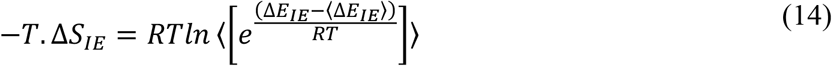

where *R* is the ideal gas constant, and Δ*E*_*IE*_ represents the interaction energy defined as the sum of Coulombic and Lennard–Jones contributions. The angle brackets ⟨⟩ denote ensemble averaging over configurations sampled from molecular dynamics trajectories.

The interaction entropy (IE) method has been further generalized in subsequent studies within a more rigorous statistical framework. In this formulation, the free energy expression derived from the exponential averaging in the IE method can be expanded as:

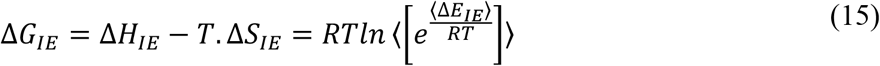

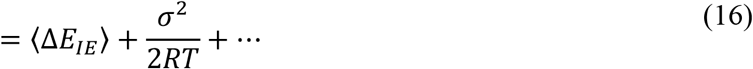

Using a second-order cumulant expansion (C2 approximation), the entropic contribution can be simplified to:

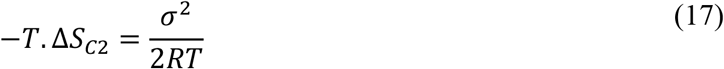

where σ^2^ corresponds to the variance of the interaction energy fluctuations. This term can be readily obtained from the fluctuations of the interaction energy along the MD trajectory.

The standard MM-PBSA method is obtained the binding free energy by summing individual energetic contributions. This approach provides a favorable compromise between computational cost and accuracy. However, it is still an approximate method, and thus standard MM-PBSA protocols often mis-predict BFEs. In literature, several studies have extended this additive framework by introducing empirical scaling of individual energy components through linear regression or fitting procedures against experimental data, aiming to improve predictive performance.

Recently, Zhang et al. [22,23] introduced a novel energy estimation approach, termed PBSA_E, which is defined as follows:

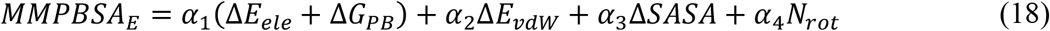

The coefficients α_1_, α_2_, α_3_, and α_4_ are empirical parameters determined through least-squares fitting against corresponding experimental binding affinities.

The coefficients *α*_1_, *α*_2_, *α*_3_ and *α*_4_ are empirical parameters determined through least-squares fitting against corresponding experimental binding affinities. In this formulation, *N*_*rot*_ represents the number of rotatable bonds of the ligand and it is used as a simple descriptor to account for entropic contribution with ligand binding.

Akkus et al. [20] further expanded this formulation by introducing fully empirical parameters for each energy component, expressing it as follows:

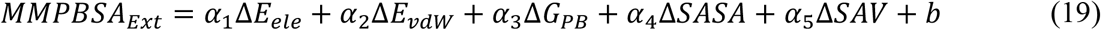

*α*_1_, *α*_2_, *α*_3_, *α*_4_, *α*_5_ coefficients and b are fitting parameters obtained using different approaches (SASA/ASA + SAV/PCAV/POAV) for the nonpolar term.

Huang et al. [19] proposed four fitted free energy estimators (equations 20-23) that integrate MM-PBSA-derived enthalpic components with interaction entropy (IE) corrections to account for entropic effects. These models were constructed using multivariate linear regression, where individual energy terms were systematically weighted to improve agreement with experimental binding affinities.

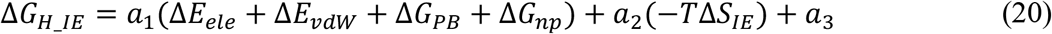

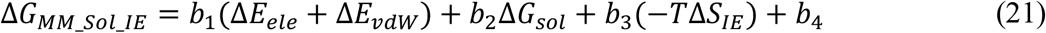

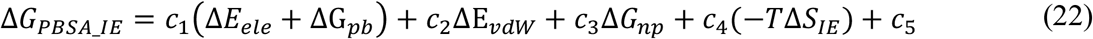

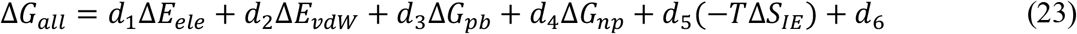

The coefficients *a*_*1*_*-a*_*3*_, *b*_*1*_*-b*_*4*_, *c*_*1*_*-c*_*5*_ and *d*_*1*_*-d*_*6*_ are determined through least-squares fitting against the corresponding experimental binding affinities.

While regression-based reweighting approaches often improve the apparent agreement between predicted and experimental binding affinities, they implicitly assume that the energy components provide independent information. However, this assumption may not be valid for MM-PB(GB)SA energy decomposition terms, as many of them come from related physical processes and tend to be statistically dependent. When strongly correlated variables are included simultaneously in a regression model, coefficient estimates can become unstable, their physical interpretation may be obscured, and the resulting models may be sensitive to overfitting. Consequently, understanding the degree of collinearity among MM-PB(GB)SA energy terms is essential for assessing the validity, robustness, and transferability of regression-based binding free energy models.

Herein, we systematically examine the statistical dependencies underlying MM-PB(GB)SA energy decomposition and their impact on binding free energy (BFE) prediction in protein–peptide complexes. Our results reveal that key energy components are strongly interdependent rather than independent as commonly assumed. Specifically, Coulomb interaction energies and polar solvation terms (PB/GB) exhibit near-perfect collinearity, consistent with their shared physical origin as vacuum electrostatics modulated by solvent screening. Similarly, van der Waals interactions show strong correlation with nonpolar solvation contributions, both largely governed by buried surface area. These trends are consistently observed across both single-trajectory and three-trajectory protocols. Overall, these findings highlight a fundamental limitation of treating decomposed MM-PB(GB)SA energy terms as independent variables in regression-based models and instead support the use of only one of the strongly correlated terms or aggregated polar, nonpolar, and entropic contributions to achieve more robust, interpretable, and transferable predictions of binding free energy.

From a strict mathematical perspective, any multivariable function can be defined as;

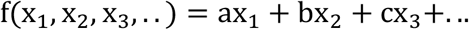

if there is a strong proportionality between any of the variables (i.e., one is dependent variable) such as;

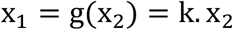

the original function can be reduced to;

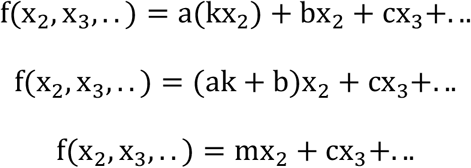

or equivalently,

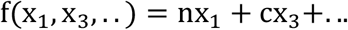

but not,

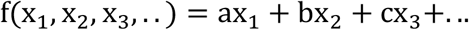

The result is that a function needs to be defined by pure independent variables in order to apply multivariant linear regression fit. Otherwise, it would just be overfitting.

Furthermore, to preserve the physical contribution of both variables without overfitting, the function can be expressed as;

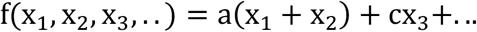

assuming 1:1 proportionality between the first two variables. If they are inversely proportional, the function can also be given by;

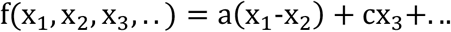

Based on collinearity results, we show MMP(G)BSA model can be at most expanded to as follows:

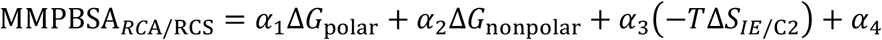

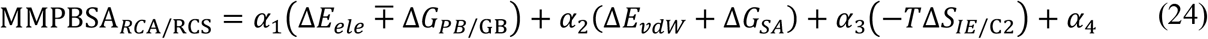

where MMPBSA_*RCA*/RCS_ represents the MMPBSA predicts free energy change which is reduced by collinearity analysis in the additive (RCA, Δ*G*_polar_ = Δ*E*_*ele*_ + Δ*G*_*PB*/GB_) or subtractive modes (RCS, Δ*G*_polar_ = Δ*E*_*ele*_-Δ*G*_*PB*/GB_). Note that the negative sign in RCS mode is due to inverse correlation of electrostatic term with PB/GB.

Furthermore, these findings clearly highlight the necessity of evaluating potential collinearity among decomposed MM-PB/GBSA energy terms when they are treated as independent variables in regression-based models.

## 3. Computational Methods

Custom Python scripts, along with PyMOL,[39] Openbabel[40] and Matplotlib Python library[41] were extensively used for calculations, file conversions, conformer generations, the construction and visualization and structural analyses and graphical representations.

### 3.1 System Preparation

To construct a protein-peptide (PP) dataset for binding free energy calculations, approximately 2,800 protein–protein complexes with reported K_d_ values were selected from the PDBBind database, and their crystal structures were retrieved from the Protein Data Bank (PDB). These complexes were then subjected to a series of filtering steps. First, complexes containing more than two chains were excluded. Next, metalloenzymes containing metal ions such as Zn, Ca, Cu, Fe, Co, Mg, and Mn were removed. Among the remaining two-chain complexes, structures in which the second chain contained more than 200 residues were excluded, as the first chain was defined as the protein and the second as the peptide. Finally, from the remaining complexes, structures containing missing amino acids were removed, except for residues located in the N- and C-terminal regions. Although some peptides are also proteins, we will refer to specify shorter length structure as peptides throughout the manuscript. All these filtering steps were automated using a custom python script, resulting in a final PP dataset of 138 complexes (TableS1).

### 3.2. Molecular Dynamics Simulations

All the classical MD simulations were carried out with GROMACS 2024.1 software package [42] with the Amber ff99SB-ILDN [43–45] all-atom force field implemented in Gromacs. PP complexes were placed at the center of a dodecahedral simulation box and solvated using the TIP3P water model [46], maintaining a minimum distance of 1.0 nm between the solute and the box boundaries. The systems were neutralized and salted by adding 0.15 M NaCl ions.

Energy minimization of the system was performed using two-step protocol combining the steepest descent and L-BFGS algorithms, each extending up to 50,000 steps when required. The minimization was carried out until the maximum force acting on any atom reached below 100 kJ mol^-1^ nm^-1^ using the Verlet cutoff scheme. Positional restraints were applied to the heavy atoms of the complexes during energy minimization, allowing the solvents and hydrogen atoms to relax freely.

Short-range nonbonded interactions, including both electrostatic (Coulomb) and van der Waals interactions, were calculated using a cutoff distance of 1.2 nm. Long-range van der Waals interactions used group scheme while electrostatic interactions were treated using the Particle Mesh Ewald (PME) method with a real-space cutoff of 1.2 nm, a fourth-order spline interpolation [47] and a Fourier grid spacing of 0.16 nm. The neighbor list was updated every 20 integration steps.

Following the energy minimization, each system was equilibrated using a two-step protocol consisting of NVT and NPT ensembles. Initially, a 1 ns NVT equilibration was performed to stabilize the system temperature at 310 K. The velocity-rescale (V-rescale) thermostat was applied with separate coupling groups for the complex and solvent. During this step, positional restraints were maintained on the heavy atoms of the PP complex.

Subsequently, the systems were subjected to pressure equilibration under an NPT ensemble. The pressure was first stabilized using a short 200 ps equilibration with the Berendsen barostat [48]. This was followed by a 1 ns equilibration using the Parrinello–Rahman barostat [49], maintaining the system at 1 atm. The temperature was continuously controlled at 310 K using the V-rescale thermostat throughout the equilibration phases.

After equilibration, all positional restraints were removed and production molecular dynamics simulations were carried out for 10 ns under the NPT ensemble at 310 K and 1 atm, using the V-rescale thermostat and the Parrinello–Rahman barostat. Periodic boundary conditions were applied in all directions. The leap-frog integrator (md) was used with a 2-fs time step. All bonds involving hydrogen atoms were constrained using the LINCS algorithm [50]. Trajectory snapshots were saved every 10 ps, resulting in 1000 frames over the 10 ns simulation. Five independent replicas were performed to ensure statistical reliability of the results.

Since both single-trajectory (ST) and three-trajectory (3A) approaches were employed, MD simulations were performed for the complex as well as for the individual chains under identical simulation conditions.

### 3.3. MM-PB(GB)SA Free Energy Calculations

Binding free energy calculations were performed using the gmx_MMPBSA [51] by employing both single-trajectory and three-trajectory approaches. A total of 1000 frames extracted from the MD simulations were used for all calculations. The binding free energy was decomposed into molecular mechanics, solvation, and entropic contributions using the gmx_MMPBSA framework. The molecular mechanics term (electrostatic and van der Waals interactions) was calculated using the ff99SB force field. Polar solvation effects were estimated using Poisson–Boltzmann (PB) and Generalized Born (GB) implicit solvent models, while the non-polar contribution was approximated using the solvent-accessible surface area (SASA) only model. The internal dielectric constant of both the protein and peptide was set to ε_in_ = 1.0. For PB calculations, the ionic strength was set to 0.150 M, while for GB calculations, the igb = 2 model [52] was used with a salt concentration of 0.150 M. The entropic contributions were estimated for single trajectory approach using interaction entropy (IE) [38], which estimates entropy from fluctuations in the interaction energy along the MD trajectory and the C2 entropy [53] method which provides an alternative statistical approximation of configurational entropy based on higher-order correlation of energy fluctuations implemented in gmx_MMPBSA.

### 3.4. Correlation Analysis and Principal Component Analysis of Energy Components

To evaluate statistical dependencies among MM-PB(GB)SA energy components, pairwise Pearson correlation analysis was performed using ensemble-averaged values obtained from multiple MD trajectories and independent replicas. This analysis was used as an exploratory tool to assess linear relationships among electrostatic, van der Waals, and solvation energy terms, as well as entropy-related descriptors.

To further investigate the intrinsic dimensionality of the enthalpic energy landscape, principal component analysis (PCA) was applied to MM-PB(GB)SA energy components (ΔE_ELE_, ΔE_vdW_, ΔG_GB/PB_, ΔG_NPOLAR/SURF_). All variables were standardized to zero mean and unit variance prior to PCA to eliminate scale-dependent effects. PCA was used to identify dominant modes of variance, characterize covariance patterns among energetic descriptors, and evaluate whether the nominal polar and nonpolar energy groupings correspond to statistically independent dimensions. Pearson correlation coefficients were additionally calculated between principal component scores and experimental binding free energies.

Entropy-related descriptors (interaction entropy, IE, and configurational entropy, C2) were excluded from PCA because they are not additive components of the MM-PB(GB)SA energy decomposition, but fluctuation-derived observables obtained from molecular dynamics trajectories. Consequently, PCA was restricted to the enthalpic energy terms, whereas entropy descriptors were analyzed separately.

All analyses were performed using Python-based scientific libraries.

## 4. Results and Discussion

We thoroughly evaluated the performance of popular end-state free energy methods, MM-PBSA and MM-GBSA, for protein–protein complexes using both single trajectory (ST), multi-trajectory (3A) approaches. To characterize the statistical relationships among decomposed energy terms, pairwise correlation analysis, principal component analysis (PCA), and frame-resolved trajectory analysis were performed. These analyses were designed not only to assess energetic trends associated with binding but also to investigate the intrinsic dependency structure and dimensionality of MM-PB(GB)SA energy decomposition.

### 4.1. Correlation of energy components across systems

To investigate how individual MM-PB(GB)SA energy components relate to each other across the dataset, we first analyzed pairwise correlations between all terms using the single-trajectory protocol (**Figure 2**). All energy terms are reported as ensemble-average quantities obtained over both MD trajectory frames and independent replicas. Throughout this work, such averages are denoted as ⟨ΔE⟩.

**Figure 2.**
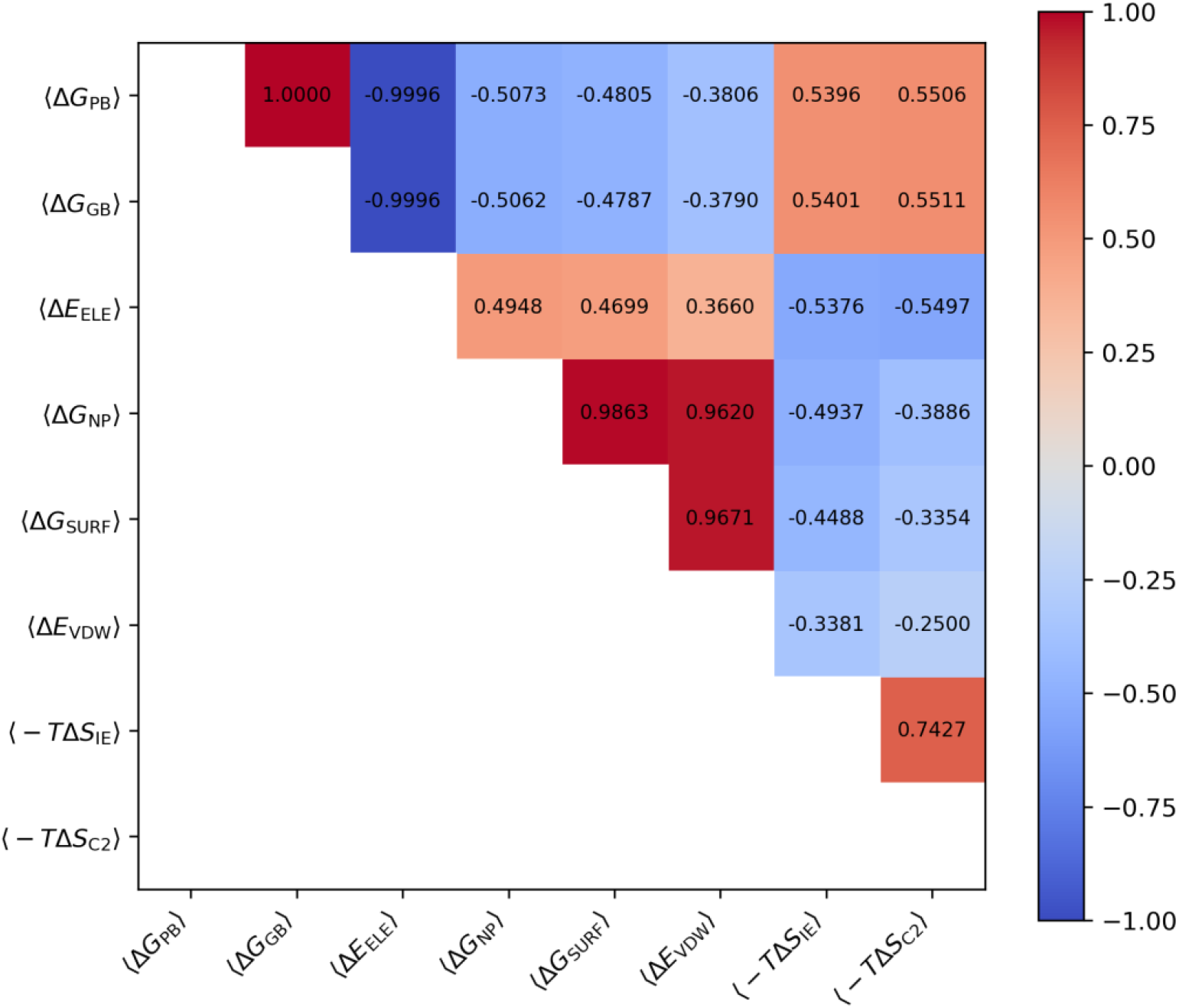
Pairwise correlation (Pearson, R) heatmap of MM-PB(GB)SA energy components across all protein-protein complexes based on the single-trajectory protocol. Strong correlations between electrostatic and polar solvation terms, as well as between van der Waals and nonpolar contributions, indicate pronounced multicollinearity within the MM-PB(GB)SA energy decomposition framework.

The correlation matrix reveals a pronounced degree of linear dependence among several energetic components, indicating that MM-PB(GB)SA decomposition terms are not statistically independent descriptors but rather partially coupled representations of underlying physical interactions. The energy terms are organized into two dominant energetic groups corresponding to polar and nonpolar contributions.

A near-perfect positive correlation was observed between the polar solvation terms ΔG_PB_ and ΔG_GB_ (Pearson R = 1.00), indicating that the Poisson–Boltzmann and Generalized Born models capture highly similar energetic trends across the present dataset, despite differences in their underlying electrostatic formulations and computational performance. Although absolute energy values may differ between the two approaches, both models provide nearly equivalent relative descriptions of polar solvation behavior.

In contrast, both polar solvation terms exhibit an almost perfect inverse correlation with the electrostatic interaction energy ΔE_ELE_ (Pearson R = −0.999), demonstrating a strong electrostatic compensation effect between gas-phase Coulombic interactions and solvent screening contributions. This is consistent with the physical formulation of these terms, where polar solvation effectively acts as a screening correction to vacuum electrostatics. Similar compensation behavior has also been described in the MM/PBSA_E_ framework reported in literature, where electrostatic and polar solvation contributions are treated as a combined energetic descriptor [22]. Consequently, assigning independent empirical fitting parameters to ΔE_ELE_ and ΔG_PB/GB_ may introduce redundancy and increase the risk of overfitting. These findings clearly indicate that electrostatic energy and polar solvation energy change in opposite directions. During complex formation, as electrostatic interactions become more favorable (i.e., more negative ΔE_ELE_), the cost of stabilizing this charge distribution of the solvent increases. This behavior represents a characteristic thermodynamic compensation effect in implicit solvent models.

A similarly strong correlation was observed between the nonpolar solvation terms ΔG_NP_ and ΔG_SURF_ (Pearson R=0.986). Both terms correspond to the change in the free energy of the solvation, which is the change in nonpolar interactions upon binding. The high correlation between these terms is understandable because they appear in two different equations and obtained from solvent accessible surface area (SASA) with an empirical proportionality parameter, γ which corresponds to surface tension coefficient. It is known that a correlation slightly lower than 1 is due to changes in the probe radius and the empirical scaling parameters used.

Interestingly, van der Waals interactions (ΔE_vdW_) also exhibit strong positive correlations with the nonpolar solvation terms (ΔG_NP_, ΔG_SURF_) (R = 0.96–0.97). Although ΔE_vdW_ is calculated from intermolecular Lennard–Jones interactions and the nonpolar solvation terms are derived from solvent-accessible surface area (SASA) based models, both quantities are fundamentally governed by intermolecular contact formation and interface burial. This observation suggests that these terms may not provide fully independent information within empirical fitting procedures, as both largely reflect the extent of hydrophobic packing and interface formation.

By contrast, electrostatic interactions (ΔE_ELE_) display only moderate correlations with the nonpolar solvation terms ΔG_NP_ and ΔG_SURF_ (R = 0.47–0.49), indicating that polar and nonpolar energetic contributions remain comparatively decoupled and can be considered largely independent energetic dimensions within the present dataset.

Entropy-related descriptors derived from interaction entropy (IE) and C2 entropy approaches exhibit similar correlation patterns with MM-PB(GB)SA energy components. Both descriptors show moderate positive correlations with polar solvation energies (ΔG_PB/GB_) (R = 0.54–0.55) and moderate negative correlations with electrostatic interaction energies (ΔE_ELE_) (R = −0.54 to −0.55). In addition, IE and C2 are somewhat strongly correlated with each other (R = 0.74), indicating that they capture partially overlapping thermodynamic information despite differences in theoretical formulation and computational implementation. In practice, these approaches are generally treated as alternative entropy estimators, and only one of them is optionally included in the energy formulation.

Collectively, these findings demonstrate that MM-PB(GB)SA energy terms exhibit substantial multicollinearity and cannot be regarded as fully independent energetic descriptors. Several energetic components therefore contain partially redundant information, which may reduce the interpretability and robustness of statistical models that assume independent predictors. Importantly, these correlation patterns are consistently preserved across both single-trajectory and three-trajectory protocols (**Figure S1**) indicating that the observed multicollinearity is intrinsic to the MM-PB(GB)SA decomposition framework rather than an artifact of a specific simulation protocol. This behavior is consistent with the non-orthogonal nature of energy decomposition schemes in end-state free energy methods, where individual terms are defined according to distinct physical models rather than statistically independent variables.

The observed dependency structure can be rationalized based on the physical definitions of the MM-PB(GB)SA energy terms. Electrostatic interactions (*E*_ELE_) are evaluated in a theoretical “vacuum” or “gas phase,” while solvation terms (*G*_GB/PB_) account for the dielectric screening effects that oppose these Coulombic forces. Crucially, this “gas phase” energy does not reflect conformations sampled in an actual vacuum. Instead, it is calculated by stripping explicit solvent molecules from an aqueous trajectory, meaning the underlying structural ensemble retains solvent-induced conformational characteristics. Similarly, van der Waals interactions (*E*_vdW_) primarily characterize intermolecular packing and dispersion forces, whereas nonpolar solvation terms (*G*_SA_) scale with buried solvent-accessible surface area (SASA). Consequently, the energy decomposition naturally organizes into two dominant energetic blocks corresponding to polar and nonpolar interaction modes.

However, pairwise correlations alone do not fully characterize the intrinsic statistical dimensionality of MM-PB(GB)SA energy space. Although the decomposition formally separates energetic contributions into polar and nonpolar terms, it remains unclear whether these groups represent truly independent energetic dimensions or instead reflect correlated manifestations of a smaller number of underlying thermodynamic modes. To address this question, principal component analysis (PCA) was performed to identify the dominant covariance directions governing the energetic landscape.

### 4.2. Principal Component Analysis of MM-PB/GBSA Energy Components

To further investigate the covariance structure and intrinsic dimensionality of the MM-PB(GB)SA energy landscape, principal component analysis (PCA) was performed on the decomposed energetic terms. As shown in **Figure 3a**, the first principal component (PC1) accounts for the majority of the total variance in both MM-GBSA (70.8%) and MM-PBSA (71.5%) datasets, indicating that the energetic space is highly constrained along a dominant covariance axis. The second principal component (PC2) captures an additional substantial fraction of the variance (MM-GBSA: 28.5%; MM-PBSA: 27.8%), whereas higher-order components contribute negligibly (<1%). Overall, the MM-PB(GB)SA energetic landscape can therefore be effectively approximated within a low dimensional two-component subspace, reflecting strong redundancy among decomposed energetic terms.

**Figure 3.**
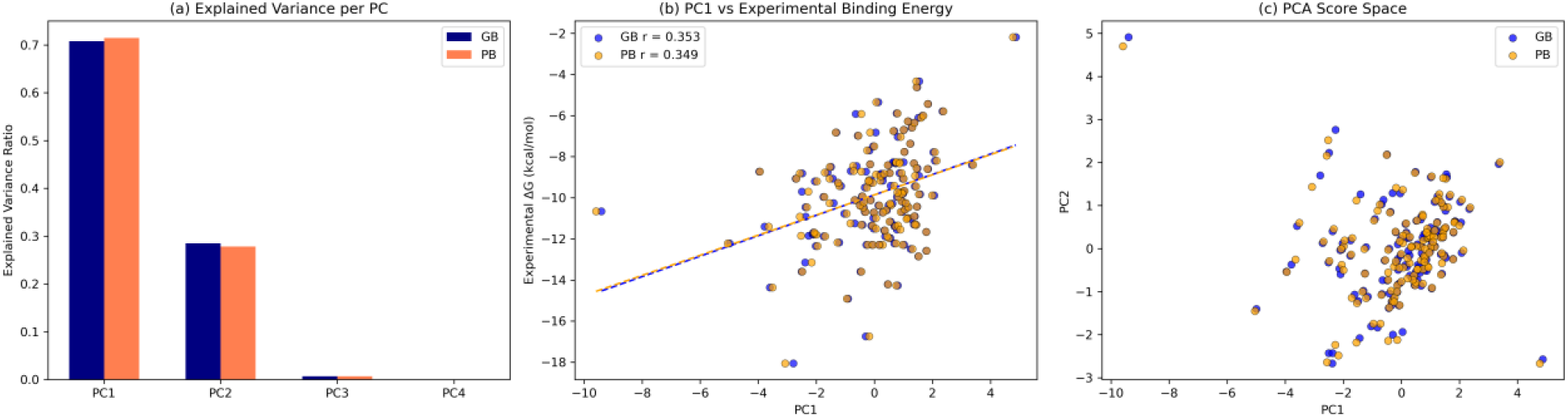
Principal component structure and experimental relevance of MM-PB(GB)SA energy space based on the ST-approach: (a) Explained variance of principal components for MM-PB(GB)SA energy terms, showing dominance of the first two components. (b) Correlation between the first principal component (PC1) and experimental binding free energies (Experimental ΔG) for MM-GBSA and MM-PBSA models, showing only a moderate relationship for both MM-GBSA and MM-PBSA. (c) Projection of complexes onto the first two principal components, revealing a continuous distribution of energetic states without clear clustering. For 3A-approach, refer to **Figure S2**.

The PCA loading structure (**Figure 4**) further clarifies the physical relationships among the energetic components. Electrostatic interaction energies and polar solvation terms project in opposite directions within PC space, consistent with the strong antagonistic compensation observed in the correlation analysis. In contrast, van der Waals interactions and nonpolar solvation terms project in similar directions, indicating coordinated behavior associated with intermolecular packing and solvent-accessible surface burial. Collectively, the loading structure reveals a dominant electrostatic–solvation compensation axis together with a coupled nonpolar interaction mode, reinforcing the presence of substantial multicollinearity within the MM-PB(GB)SA decomposition framework.

**Figure 4.**
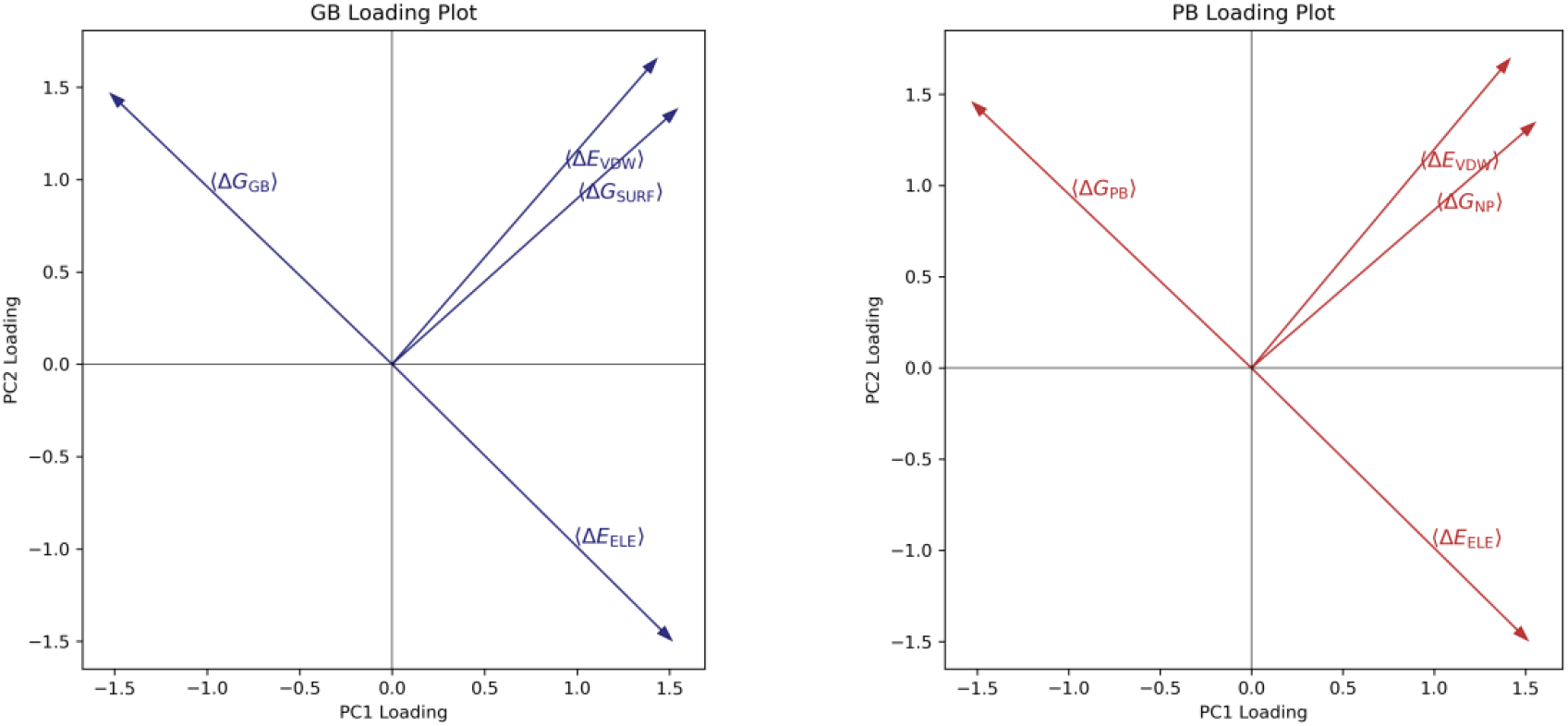
PCA loading plots of MM-GBSA and MM-PBSA energy components based on the ST-approach projected onto the PC1–PC2 space. Electrostatic and polar solvation terms project in opposite directions, indicating strong electrostatic compensation behavior, whereas van der Waals and nonpolar solvation terms project, similarly, reflecting coupled hydrophobic and surface-area–driven contributions. The overall loading structure reveals substantial multicollinearity and a dominant electrostatic–solvation covariance axis within the MM-PB(GB)SA energetic landscape. For 3A-approach, refer to **Figure S3**.

Despite explaining the majority of the variance within the energetic decomposition, PC1 exhibits only moderate correlations with experimental binding free energies for both MM-GBSA (R = 0.353) and MM-PBSA (R = 0.349) (

**Figure 3b**). This observation suggests that the dominant covariance mode primarily reflects internal energetic coupling within the decomposition scheme rather than directly encoding experimental binding affinity. The relatively weak relationship with experimental ΔG further indicates that additional thermodynamic factors not explicitly represented within the additive enthalpic decomposition may contribute significantly to binding energetics. Configurational entropy and dynamic fluctuations sampled during MD simulations are not directly represented within the PCA framework and may partially account for the observed discrepancy.

Projection of the complexes onto the first two principal components (**Figure 3c**) reveals a continuous energetic distribution rather than discrete clustering behavior, indicating gradual variation in energetic profiles across the dataset. Notably, MM-PBSA and MM-GBSA models exhibit nearly identical variance distributions and loading structures, suggesting that the overall covariance organization of the energetic landscape is largely preserved despite differences in the underlying polar solvation models.

### 4.3. Pairwise Compensation Analysis of MM-PB/GBSA Energy Components

To further elucidate the physical relationships underlying the correlation structure and principal component decomposition, pairwise linear regression analyses were performed between the highly correlated energy components for both individual replicas and their averages (**Figure 5**). Unlike correlation matrices, these pairwise relationships explicitly reveal the energetic compensation and coupling behaviors.

**Figure 5.**
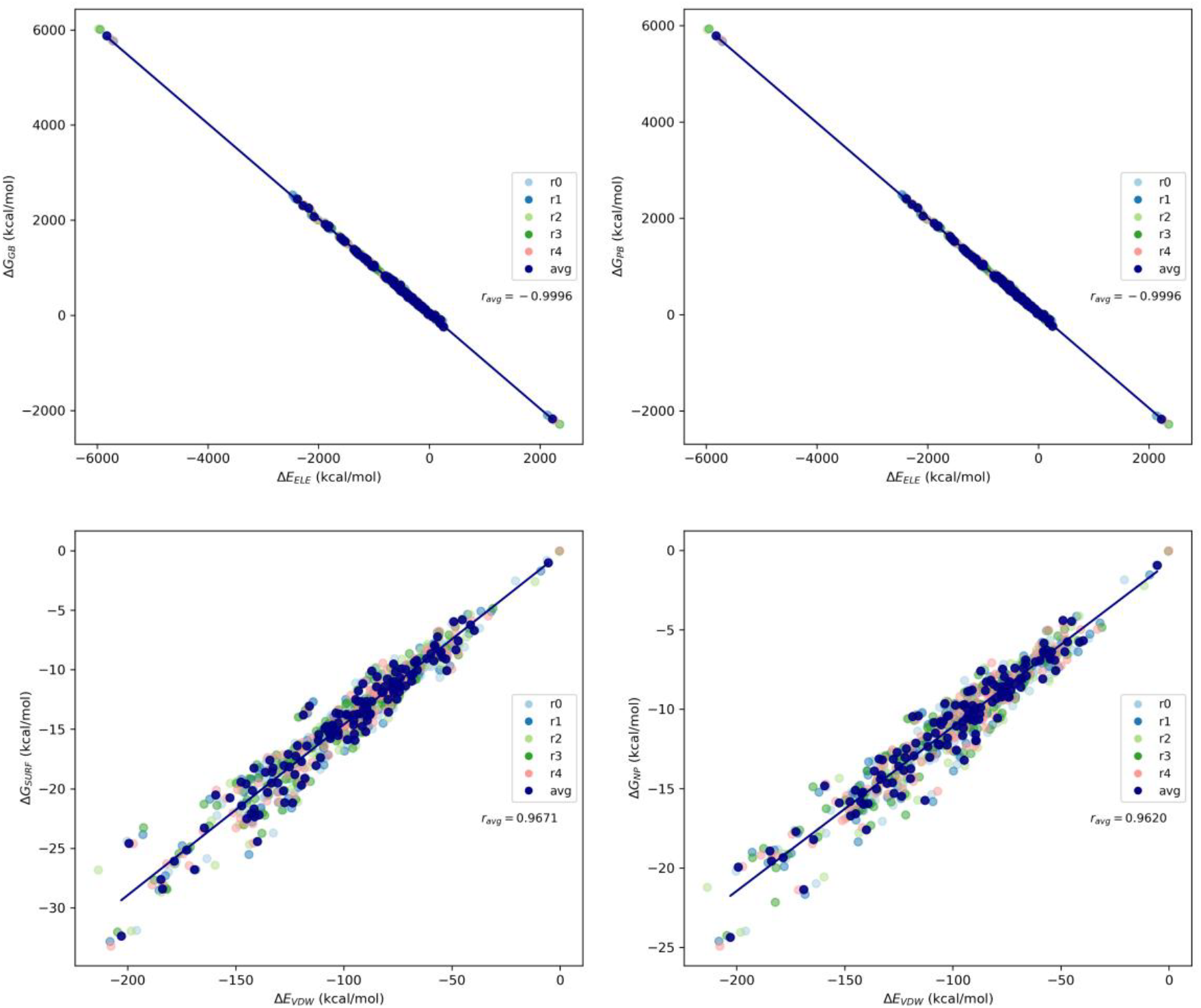
Pairwise energetic compensation relationships among MM-PB(GB)SA energy components across all protein-protein complexes based on the ST-approach (For 3A-approach, refer to **Figure S4**). Top panels show the nearly perfect inverse correlation between electrostatic interaction energy (ΔE_ELE_) and polar solvation energies (ΔG_PB/GB_), highlighting electrostatic compensation effects intrinsic to implicit solvent models. Bottom panels illustrate the strong positive coupling between van der Waals interactions (ΔE_vdW_) and nonpolar solvation contributions (ΔG_NP_ and ΔG_SURF_). Colored points correspond to independent simulation replicates (r0–r4), while the regression line represents the averaged values.

A near-perfect inverse linear relationship is observed between the electrostatic interaction energy (ΔE_ELE_) and polar solvation energies (ΔG_PB_ and ΔG_GB_), highlighting the strong coupling between these terms and the role of polar solvation as an electrostatic screening correction characteristic of implicit solvent models. This trend indicates a strong coupling between favorable Coulombic interactions and unfavorable polar solvation contributions during complex formation. This behavior is inherent to the thermodynamic formulation of MM-PB(GB)SA, in which opposes energetic contributions, such as gas-phase electrostatics and polar solvation, are explicitly separated and subsequently recombined to estimate the final binding free energy. Consequently, the calculated binding energy primarily reflects the balance between strongly coupled energetic terms rather than fully independent physical contributions.

In contrast, van der Waals interactions (ΔE_vdW_) exhibit a strong positive correlation with nonpolar solvation terms (ΔE_SURF_ and ΔE_NP_), suggesting a cooperative hydrophobic contribution to binding stabilization. This coupling indicates that dispersion-driven contacts and solvent-accessible surface effects evolve in a coordinated manner, reflecting their shared dependence on hydrophobic contact formation and solvent-accessible surface exposure. Taken together, these trends underscore the distinct but complementary roles of electrostatic and nonpolar interactions governing binding free energy landscapes, where solvent-mediated effects act as key modulators of energetic balance.

Importantly, these relationships are consistently preserved across independent simulation replicates (r0–r4), confirming the robustness of the observed energetic coupling patterns. Moreover, comparison between single-trajectory and three-trajectory protocols reveals similar correlation trends, suggesting that multicollinearity is an intrinsic feature of MM-PB(GB)SA energy decomposition rather than a method-dependent artifact.

These energetic coupling patterns are consistent with the principal component structure observed in the PCA analysis, where the dominant variance axis (PC1) reflects the combined covariance of electrostatic interactions, van der Waals forces, and opposing polar solvation contributions.

These energetic compensation patterns provide a physically consistent interpretation of the principal component structure observed in the PCA analysis, where the dominant variance axis (PC1) emerges from the coupled contributions of electrostatic interactions, van der Waals forces, and opposing polar solvation energies.

### 4.4. Frame-by-Frame Collinearity Analysis of MM-PB(GB)SA Energy Components Along MD Trajectories

The collinearity analyses performed so far have been presented using the average values of each MD trajectory and replica. To further elucidate the temporal nature of these relationships, frame-by-frame collinearity analyses were performed along the MD trajectories. Consistent with the ensemble-level results, electrostatic interaction energies and polar solvation terms remained strongly correlated throughout the simulations (**Figure 6**), reflecting the intrinsic electrostatic compensation behavior characteristic of implicit solvent models. In contrast, a pronounced correlation was not observed between van der Waals and nonpolar solvation terms at the frame level (**Figure 7**). This discrepancy suggests that the strong ensemble-level correlation primarily arises from slowly varying structural descriptors such as average interface burial and overall conformational compactness rather than direct instantaneous energetic coupling. While nonpolar solvation terms vary relatively smoothly with solvent-accessible surface area, van der Waals interactions are highly sensitive to transient local packing fluctuations and short-range atomic rearrangements. Consequently, the apparent vdW–nonpolar coupling represents a long-timescale statistical relationship rather than a strict frame-resolved physical correspondence.

**Figure 6.**
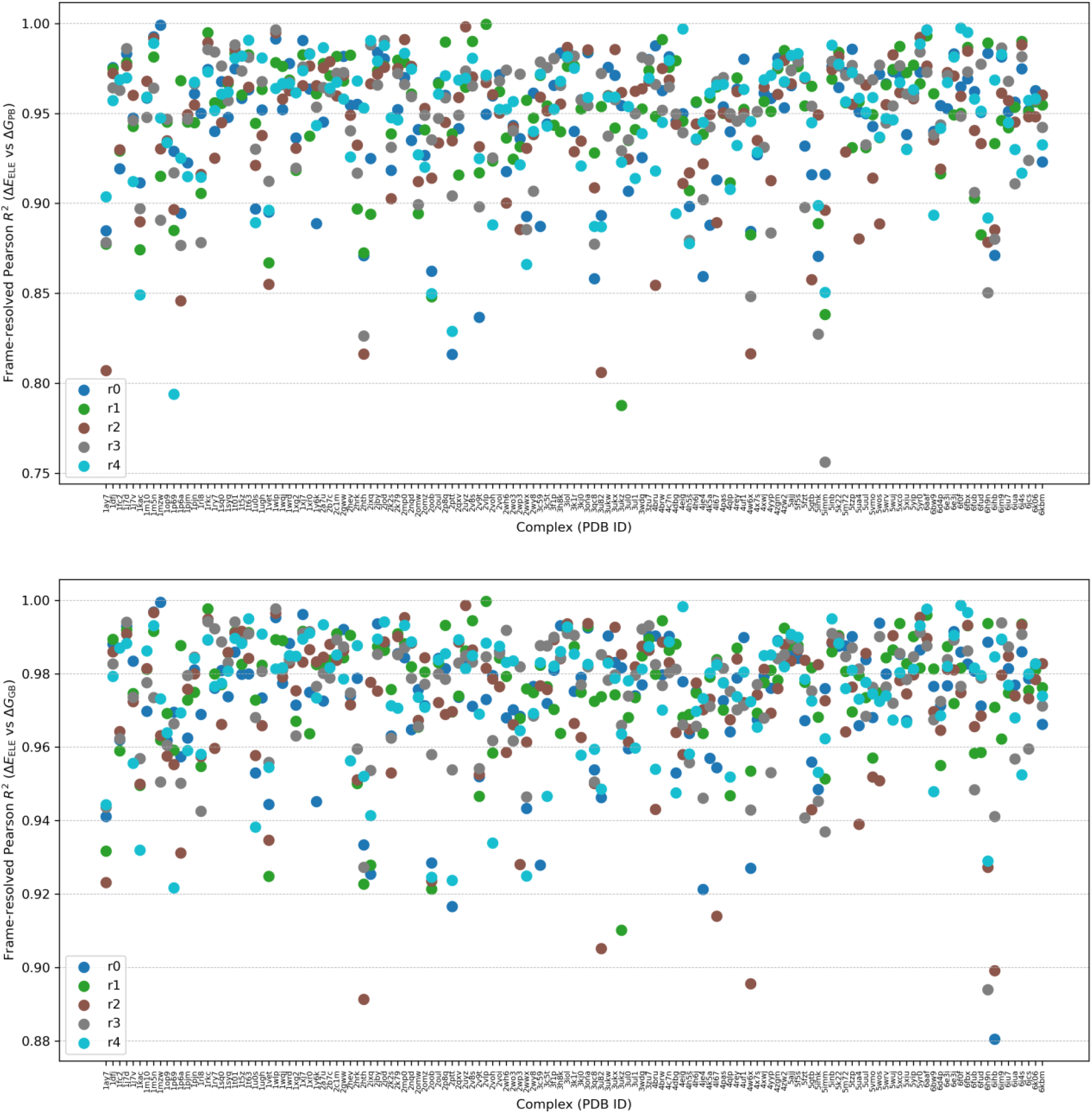
Frame-by-frame variation of the correlation between electrostatic interaction energy (ΔE_ELE_) and polar solvation energies (ΔG_PB/GB_) along the MD trajectory based on the single-trajectory protocol.

**Figure 7.**
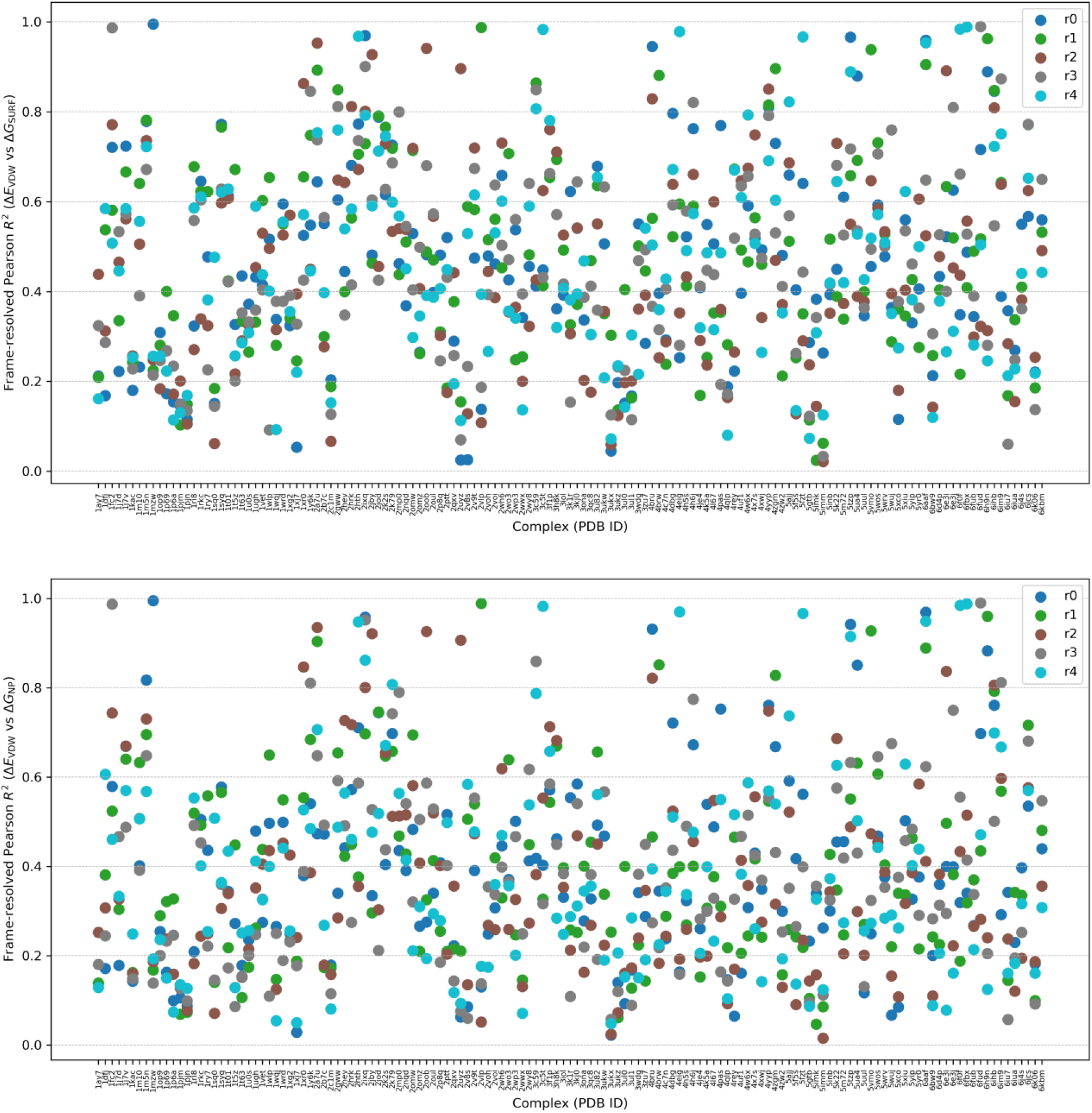
Frame-by-frame variation of the correlation between van der Waals interactions (ΔE_vdW_) and nonpolar solvation contributions (ΔG_NP_ and ΔG_SURF_) along the MD trajectory based on the single-trajectory protocol.

### 4.5. Predictive performance of different MM-PB(GB)SA approaches

To further investigate how the internal dependency structure of MM-P(G)BSA energy components influences their predictive behavior, several linear regression models previously proposed in the literature and newly introduced reduced-component (RCA/RCS) representations including IE or C2 corrections were evaluated against the experimental binding free energies (BFE). Replica-averaged energy terms obtained from the five independent MD simulations for each complex were used in all regression analyses to minimize trajectory-dependent fluctuations and emphasize ensemble-averaged energetic trends. As a reference, the original MM-P(G)BSA formulation based on the direct summation of energetic contributions was first considered:

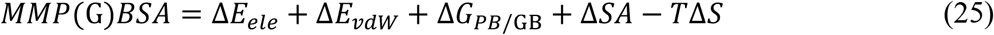

In this conventional representation, all energetic terms contribute equally with fixed coefficients, assuming that each decomposed energy component contributes equally to the total binding free energy. This formulation showed relatively weak correlations with the experimental BFEs (**Table 1**, the corresponding plots are in **Figure S5-S6**). The original MMPBSA model yielded a Pearson correlation coefficient of 0.30, while the MMGBSA model produced a slightly lower correlation of 0.27. Inclusion of entropic corrections using either the interaction entropy (IE) or C2 approaches did not improve the predictive performance. On the contrary, the correlations decreased substantially, in some cases becoming negative.

**Table 1.** Pearson correlation coefficients obtained for different MM-PBSA and MM-GBSA (values in parentheses) based regression models.

| Approach | Pearson, R | IE entropy included | C2 entropy included | Equation involved |
| --- | --- | --- | --- | --- |
| **MMPBSA** | 0.30 (0.27) | 0.10 (0.06) | -0.07 (-0.07) | 30 |
| **MMPBSA<sub>E</sub>** | 0.31 (0.28) | 0.31 (0.29) | 0.31 (0.28) | 31 |
| **MMPBSA<sub>Ext</sub>** | 0.38 (0.37) | 0.38 (0.37) | 0.39 (0.39) | 32 |
| **MMPBSA<sub>RCA</sub>** | 0.30 (0.28) | 0.3 (0.29) | 0.31 (0.28) | 33 |
| **MMPBSA<sub>RCS</sub>** | 0.36 (0.36) | 0.36 (0.36) | 0.38 (0.38) | 34 |

These observations indicate that the direct additive combination of electrostatic, van der Waals, polar solvation, nonpolar solvation, and entropy terms does not effectively reproduce the experimental energetic trends for the analyzed protein-protein complexes. The decline observed after the inclusion of entropy further demonstrates that the estimated entropy terms introduce significant statistical noise and sampling-dependent fluctuations into the models. This behavior may reflect the limited convergence and strong trajectory dependence of approximate entropy estimation methods, particularly for large and flexible protein–protein interfaces.

To investigate whether the predictive limitations originate from the predefined equal weighting of the energetic components (conventional MMPBSA), multiple linear regression models with independently optimized coefficients were constructed. The first reweighted model MMGBSA_E_ proposed by Zhang [19] grouped the electrostatic and polar solvation terms into a combined electrostatic-desolvation component:

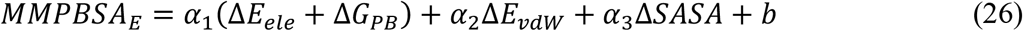

This formulation is motivated by the strong electrostatic-solvation compensation frequently observed in MM-PB(GB)SA calculations, where favorable electrostatic interactions are often counterbalanced by unfavorable polar desolvation penalties. Combining these highly correlated terms into a single descriptor reduced redundancy in the regression space and resulted in a clear improvement in the prediction of protein-ligand binding free energy. In their studies for protein-ligand systems, the Pearson correlation coefficient increased from 0.46 using the conventional MM-PBSA to 0.72 using this formulation which also included the entropic contribution [19].

In contrast, the present protein–protein study does not exhibit a similar improvement. The MMPBSA_E_ and MMGBSA_E_ formulations yield only bare improvements relative to conventional MM-PBSA, with Pearson correlation coefficients of approximately 0.31–0.28. While reweighting strategies have proven effective for protein–ligand binding affinity prediction, the improvement observed in those systems was not reproduced for the protein–protein complexes examined in this work, suggesting that the underlying energetic relationships may differ substantially between the two classes of biomolecular interactions.

The fitted coefficients revealed that the combined electrostatic-solvation contribution received a small effective weight, whereas the van der Waals contribution retained a larger coefficient. This behavior strongly suggests that electrostatic energy and polar solvation energy compensate each other. Since these terms are strongly anti-correlated and typically exhibit opposite signs, their sum significantly suppresses the informative variance available for regression. Consequently, despite the individually large magnitudes of the component terms, the net electrostatic-desolvation contribution behaves as a weak predictor.

Subsequently, another study for protein-ligand interactions [54] used a more detailed decomposition strategy in the MMPBSA_Ext_ formulation, where all energetic components are considered independently within the regression framework:

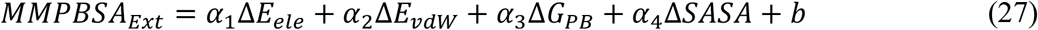

Unlike the MMPBSA_E_ representation, this formulation preserves the full energetic decomposition and allows the regression model to independently optimize the contribution of each energetic component. The study showed a substantial enhancement in predictive performance, where the Pearson correlation coefficient improved from approximately 0.42–0.47 with standard MM-PBSA to 0.85–0.88 using the MMPBSA_E_ approach [20].

The independently weighted model yielded the highest correlations observed in this study for protein-protein interactions, reaching Pearson coefficients of 0.38–0.37. Notably, the internal dielectric constant, ε_in,_ was not varied in these analyses, since a change in ε_in_ rescales the electrostatic and polar-solvation terms by a global factor that is absorbed into the fitted coefficients and therefore cannot alter the correlation of the independently weighted model.

Motivated by the strong collinearity patterns observed throughout the present analyses, we introduce a reduced-component additive (RCA) linear regression representation termed MMPBSA_RCA_. In this formulation, the electrostatic and polar solvation terms are combined into a single descriptor, while the van der Waals and nonpolar solvation terms are combined into a second descriptor:

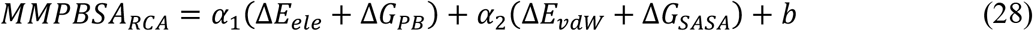

Compared to the previous formulations, MMPBSA_RCA_ explicitly reduces the dimensionality of regression space by combining pairs of highly correlated energetic terms into two physically interpretable collective descriptors. This reduced representation was designed to minimize redundancy while preserving the dominant energetic compensation behavior identified in the correlation and PCA analysis.

Despite reducing the dimensionality of the energetic space, the resulting RCA models showed similar prediction performance to the original formulation. This result suggests that a substantial fraction of the observed variance can be represented using only two dominant interaction coordinates: an electrostatic-desolvation mode and a hydrophobic packing mode. This observation is fully consistent with the PCA and collinearity analyses presented earlier, where strong dependencies were observed among electrostatic and solvation-related terms.

While the RCA formulation captures the net effect of electrostatic–solvation compensation through addition of the two terms, it does not explicitly assess the magnitude of the residual imbalance between them. To explore whether this residual component contains additional predictive information, we introduced an alternative reduced-component subtractive (RCS) model termed MMPBSA_RCS_:

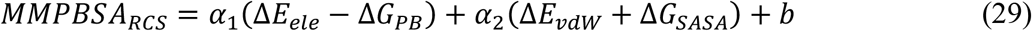

Unlike the additive RCA representation, the RCS formulation replaces the sum of the electrostatic and polar solvation terms with their difference. Since these components frequently compensate each other, the subtraction-based descriptor emphasizes the residual energetic contribution that remains after compensation and may reduce redundancy arising from their strong correlation.

The most notable improvement among the reduced-coordinate models was obtained using the RCS formulation. This model achieved correlations up to 0.38, approaching the performance of the fully independent regression model (MMPBSA_Ext_) despite using substantially fewer effective variables. This result is physically significant. In conventional MM-PB(GB)SA formulations, the electrostatic interaction and polar solvation terms are summed directly, leading to strong energetic cancellation. In contrast, the subtraction-based representation may better represent the effective electrostatic driving force by separating attractive intermolecular electrostatics from solvent-mediated desolvation penalties. The performance of the RCS model therefore suggests that the electrostatic contribution is better represented as a competition between intermolecular electrostatic attraction and desolvation cost rather than as a simple additive quantity.

Although the regression-based models improved the correlations relative to the conventional MM-PB(GB)SA formulations, the overall predictive performance remained moderate. This limitation likely reflects multiple factors beyond energetic decomposition itself, including force-field inaccuracies, implicit solvent approximations, conformational sampling limitations, and the intrinsic complexity of protein–protein binding thermodynamics.

Collectively, these results demonstrate that MM-PB(GB)SA energy decomposition terms do not behave as statistically independent physical descriptors. Instead, the results suggest that the apparent complexity of the MM-PB(GB)SA energy terms can largely be described by a small number of strongly coupled interaction patterns, primarily involving electrostatic–desolvation compensation and cooperative hydrophobic interactions arising from coupled van der Waals and nonpolar solvation contributions. The consistency between the regression analyses, collinearity measurements, and PCA results further supports the interpretation that substantial redundancy exists among the decomposed energetic components. Consequently, reduced-coordinate representations may provide more physically meaningful and statistically robust descriptions of protein–protein binding energetics than conventional additive formulations.

## 5. Conclusions

This study systematically investigated the internal dependency structure of MM-PB(GB)SA energy decomposition terms in protein–protein complexes. Correlation analysis revealed strong linear dependencies among electrostatic and solvation-related components, while principal component analysis demonstrated that the majority of variance is captured by a limited number of latent dimensions. These results demonstrate that decomposed energy terms are not statistically independent descriptors but rather correlated representations of a reduced set of underlying physical interactions.

Although the total binding free energy is formally expressed as a sum of distinct contributions, the individual MM-PB(GB)SA terms—such as electrostatic, van der Waals, polar, and nonpolar solvation energies—are not physically separable in a strictly independent manner. Instead, they represent interdependent components of the same binding process, where favorable contributions in one term are often accompanied by compensatory effects in others. This inherent coupling leads to pronounced multicollinearity and an effectively reduced dimensionality of the energy landscape.

These correlations arise naturally from the thermodynamic and methodological structure of MM-PB(GB)SA, where energy components are computed through distinct approximations but ultimately describe interconnected physical phenomena. As a result, the standard decomposition framework introduces redundancy that is intrinsic rather than purely statistical.

Our findings therefore challenge the common assumption of independence when using MM-PB(GB)SA components in regression-based or reweighting approaches. Methods such as MMPBSA_E_, MMPBSA_Ext_, and LIE-type models, which rely on independently fitted coefficients, may be affected by unstable parameter estimation and reduced interpretability under strong collinearity.

Regression-based reweighting of MM-PB(GB)SA energy components provided additional insight into the predictive structure of the system. While independently optimized linear models (e.g., MMPBSA_Ext_) yielded modest improvements in correlation with experimental binding free energies, the overall predictive performance remained limited, with Pearson coefficients not exceeding moderate values. Reduced-coordinate representations (MMPBSA_RCA_ and MMPBSA_RCS_) achieved comparable performance despite using fewer effective degrees of freedom, further supporting the presence of strong redundancy among energy components. Notably, the best-performing models consistently indicated that electrostatic and solvation contributions do not function as independent predictors but instead behave as coupled energetic modes.

Overall, this study highlights the importance of explicitly accounting for multicollinearity when developing predictive models based on decomposed free energy terms. Feature aggregation or dimensionality reduction approaches may provide more robust and physically consistent representations for machine learning applications in binding affinity prediction.

The study also uncovers the limited predictive performance of MMP(G)BSA methods for protein-protein interactions based on a large-scale systematic evaluation.

## 6. Author Information

### 6.1. Corresponding Authors

**Abdulkadir Kocak** – Department of Chemistry, Gebze Technical University, 41400 Gebze, Kocaeli, Turkey; E-Mail; orcid.org/0000-0001-6891-6929

### 6.2. Authors

**Azize SEVIM** – Department of Physics, Gebze Technical University, 41400 Gebze, Kocaeli, Turkey; E-Mail; orcid.org/0000-0002-4689-2070

### 6.3. Notes

The authors have no relevant affiliations or financial involvement with any organization or entity with a financial interest in or financial conflict with the subject matter or materials discussed in the manuscript. This includes employment, consultancies, honoraria, stock ownership or options, expert testimony, grants, patents received or pending, or royalties.

### 6.4. Author contributions

A. Kocak conceptualized the study and designed the methodology. A. Kocak and A. Sevim prepared the python scripts. A. Sevim and A. Kocak evaluated and discussed the findings. A. Sevim and A. Kocak wrote the manuscript. A. Kocak supervised the overall study. All authors approved the manuscript.

## 7 Acknowledgements

The numerical calculations reported in this paper were partially performed at TUBITAK ULAKBIM, High Performance and Grid Computing Center (TRUBA resources). This work was supported by Scientific and Technological Research Council of Turkey─TUBITAK (Project Number: **223Z125**).

The authors used AI-assisted tools for manuscript proofreading, language editing, and the development of Python scripts for data visualization. All scientific content, data interpretation, and conclusions are the work of the authors, who take full responsibility for the accuracy and integrity of the reported results.

## 8. Supporting Information

Additional figures and tables are provided in the Supporting Information.

