## Supporting Information for "Collinearity of Decomposed Energy Terms in MM-P(G)BSA Binding Free Energy Calculations of Protein-Protein Complexes"

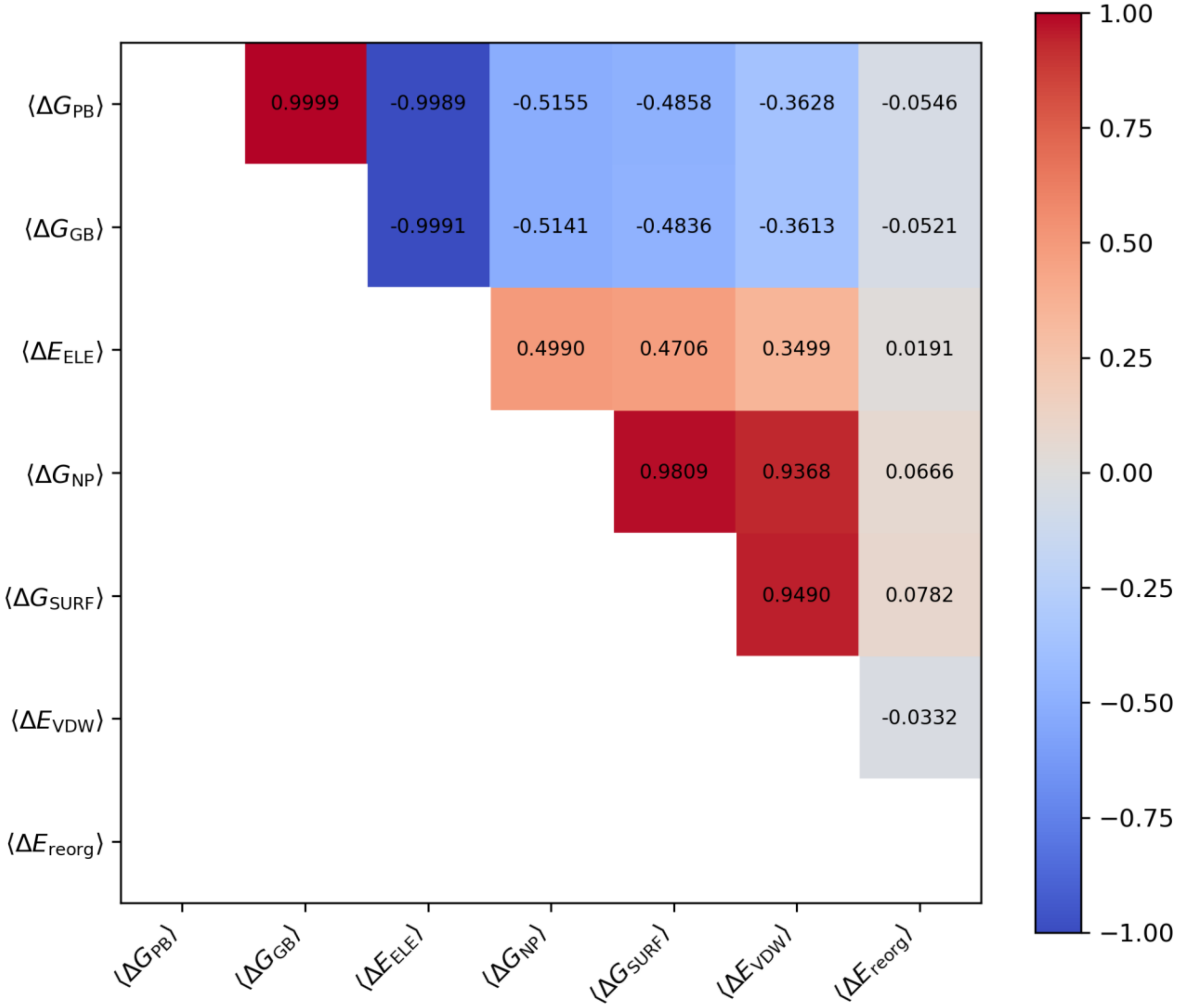


***Figure S1.*** *Pairwise correlation heatmap of MM-PB/GBSA energy components across all protein-protein complexes based on the three-trajectory protocol. Strong correlations between electrostatic and polar solvation terms, as well as between van der Waals and nonpolar contributions, indicate pronounced multicollinearity within the MM-PB(GB)SA energy decomposition.*

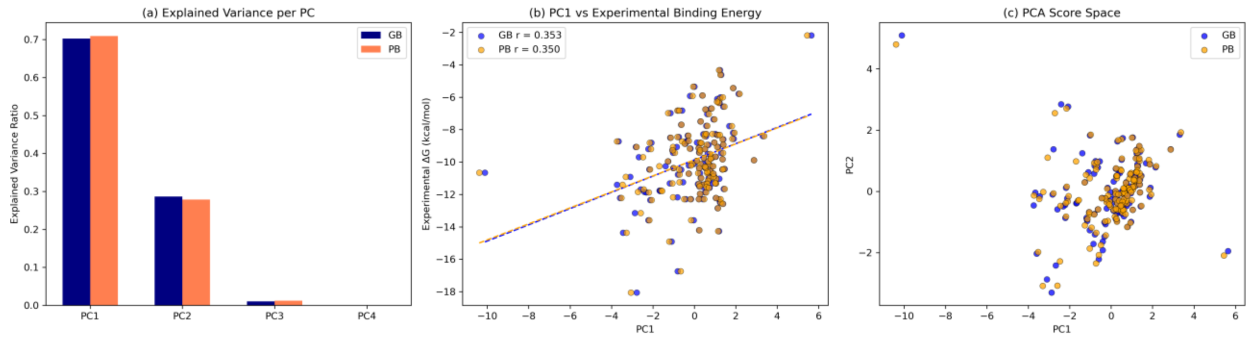

***Figure S2.*** *Principal component structure and experimental relevance of MM-PB(GB)SA energy space based on the three-trajectory protocol: (a) Explained variance of principal components for MM-PB(GB)SA energy terms, showing dominance of the first two components. (b) Correlation between the first principal component (PC1) and experimental binding free energies (Experimental ΔG) for MM-GBSA and MM-PBSA models, showing only a moderate relationship for both MM-GBSA and MM-PBSA. (c) Projection of complexes onto the first two principal components, revealing a continuous distribution of energetic states without clear clustering.*

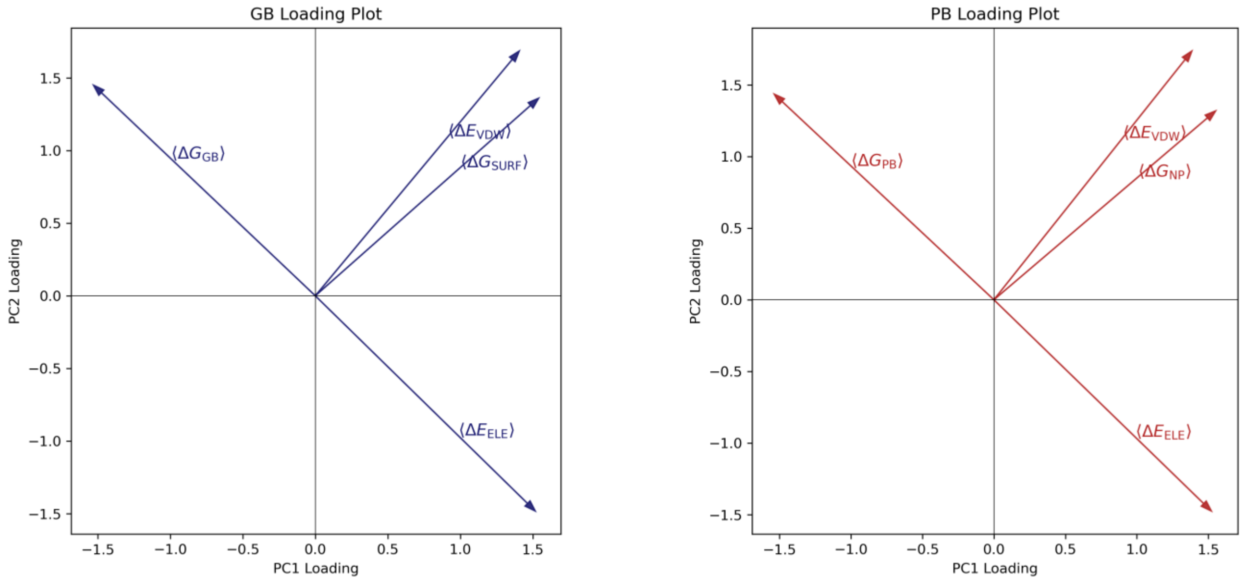


***Figure S3.*** *PCA loading plots of MM-GBSA and MM-PBSA energy components (based on the single-trajectory protocol) projected onto the PC1–PC2 space. Electrostatic and polar solvation terms project in opposite directions, indicating strong electrostatic compensation behavior, whereas van der Waals and nonpolar solvation terms project, similarly, reflecting coupled hydrophobic and surface-area–driven contributions. The overall loading structure reveals substantial multicollinearity and a dominant electrostatic–solvation covariance axis within the MM-PB(GB)SA energetic landscape.*

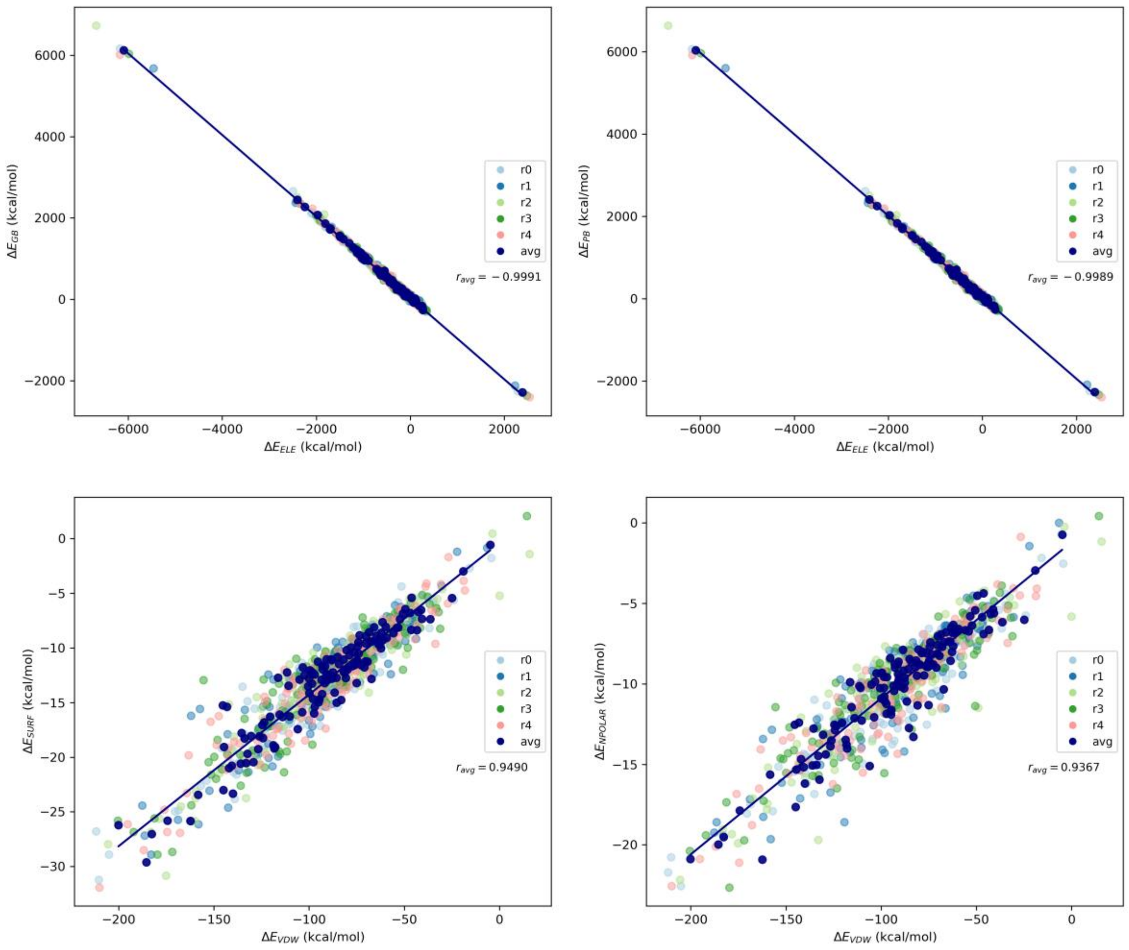


***Figure S4***. *Pairwise energetic compensation relationships among MM-PB(GB)SA energy components across all protein-protein complexes based on the three-trajectory protocol. Top panels show the nearly perfect inverse correlation between electrostatic interaction energy ($\Delta E_{ELE}$) and polar solvation energies ($\Delta G_{PB/GB}$), highlighting electrostatic compensation effects intrinsic to implicit solvent models. Bottom panels illustrate the strong positive coupling between van der Waals interactions ($\Delta E_{vdW}$) and nonpolar solvation contributions ($\Delta G_{NP}$ and $\Delta G_{SURF}$). Colored points correspond to independent simulation replicates (r0–r4), while the regression line represents the averaged values.*

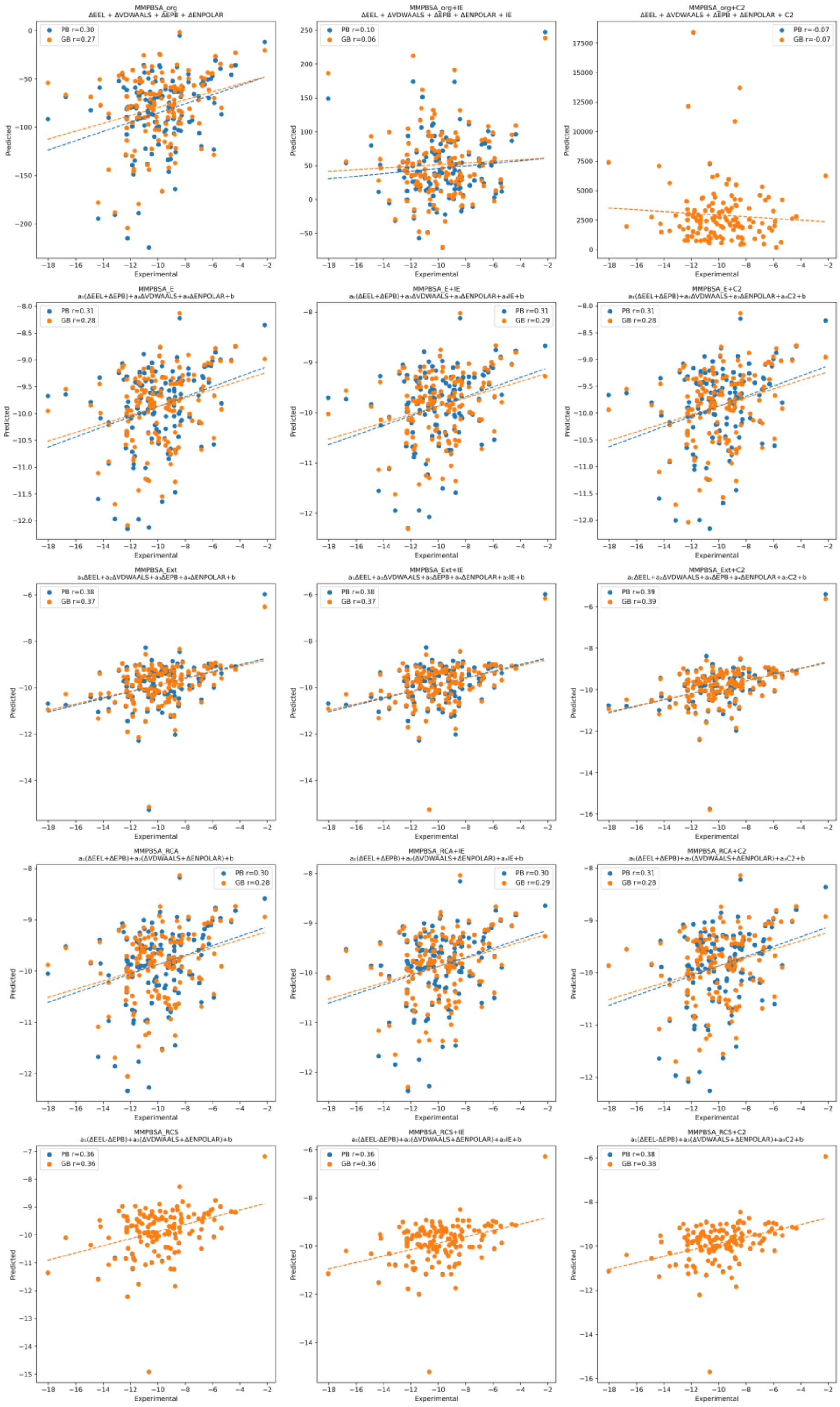


***Figure S5.*** *Comparison of experimental binding affinities with values predicted by the original MM-PB(GB)SA models, interaction entropy (IE)-corrected models, and C2 entropy-corrected models based on the single-trajectory protocol. The figure includes the MMPBSA_org, MMPBSA_E, MMPBSA_Ext, MMPBSA_RCA, and MMPBSA_RCS formulations together with their corresponding +IE and +C2 variants. Scatter plots show predicted versus experimental values, and dashed lines indicate linear fits. Pearson correlation coefficients (r) are reported for both PB- and GB-based models.*

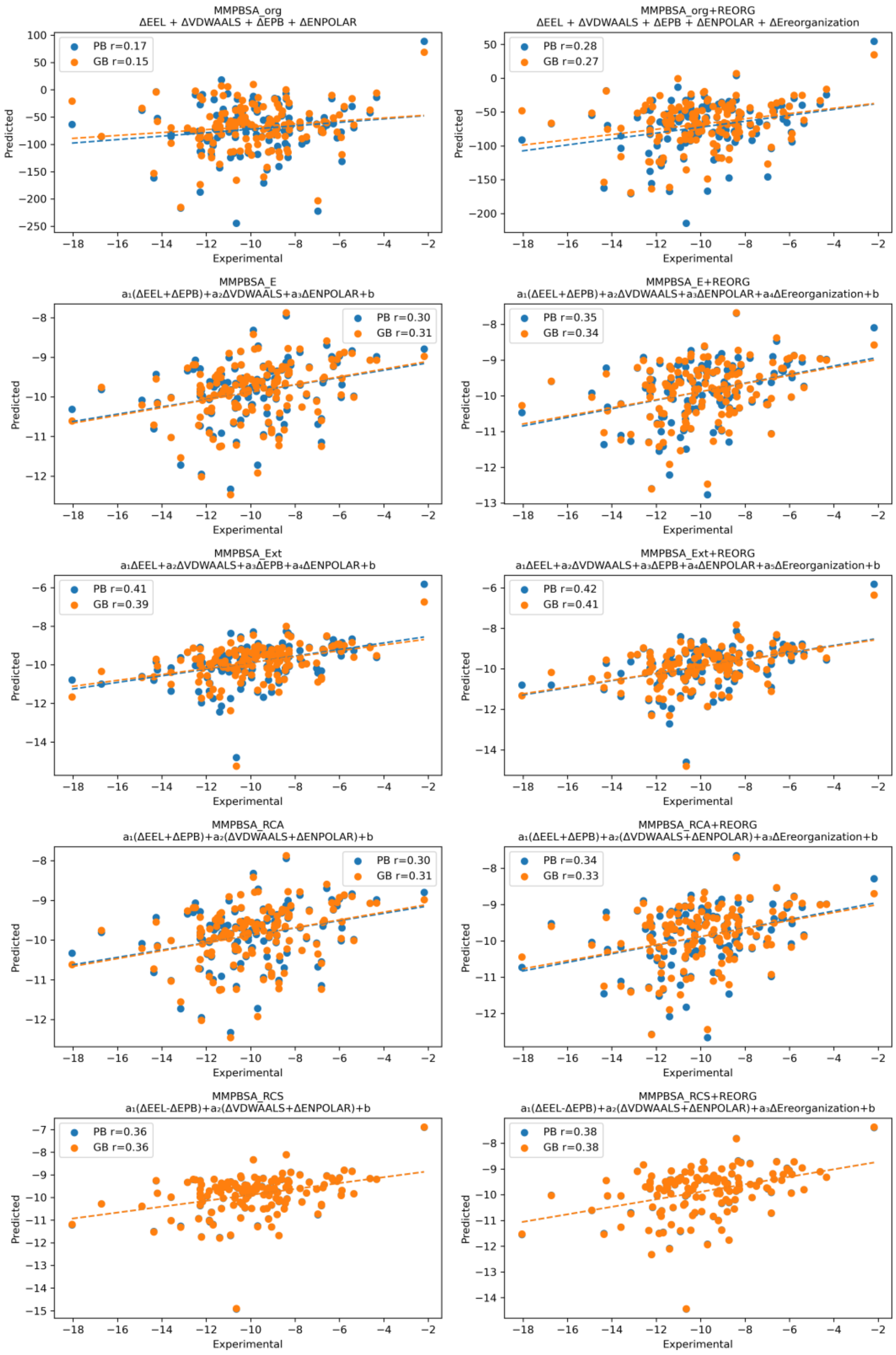


***Figure S6.*** *Comparison of experimental binding affinities with values predicted by the original MM-PB(GB)SA models based on the three-trajectory protocol. The figure includes the MMPBSA_org, MMPBSA_E, MMPBSA_Ext, MMPBSA_RCA, and MMPBSA_RCS formulations together with their corresponding ΔEreorganization variants. Scatter plots show predicted versus experimental values, and dashed lines indicate linear fits. Pearson correlation coefficients (r) are reported for both PB- and GB-based models.*
